# A strategy to detect metabolic changes induced by exposure to chemicals from large sets of condition-specific metabolic models computed with enumeration techniques

**DOI:** 10.1101/2023.06.30.547200

**Authors:** Louison Fresnais, Olivier Perin, Anne Riu, Romain Grall, Alban Ott, Bernard Fromenty, Jean-Clément Gallardo, Maximilian Stingl, Clément Frainay, Fabien Jourdan, Nathalie Poupin

## Abstract

**Background:** The growing abundance of *in vitro* omics data, coupled with the necessity to reduce animal testing in the safety assessment of chemical compounds and even eliminate it in the evaluation of cosmetics, highlights the need for adequate computational methodologies. Data from omics technologies allow the exploration of a wide range of biological processes, therefore providing a better understanding of mechanisms of action (MoA) related to chemical exposure in biological systems. However, the analysis of these large datasets remains difficult due to the complexity of modulations spanning multiple biological processes.

**Results:** To address this, we propose a strategy to reduce information overload by computing, based on transcriptomics data, a comprehensive metabolic sub-network reflecting the metabolic impact of a chemical. The proposed strategy integrates transcriptomic data to a genome scale metabolic network through enumeration of condition specific metabolic models hence translating transcriptomics data into reaction activity probabilities. Based on these results, graph algorithm is applied to retrieve user readable sub-networks reflecting the possible metabolic MoA (mMoA) of chemicals. This strategy has been implemented as a three-step workflow. The first step consists in building cell condition-specific models reflecting the metabolic impact of each exposure condition while taking into account the diversity of possible optimal solutions with a partial enumeration algorithm. In a second step, we address the challenge of analyzing thousands of enumerated conditions-specific networks by computing differentially activated reactions (DARs) between the two sets of enumerated possible condition-specific models. Finally, in the third step, DARs are grouped into clusters of functionally interconnected metabolic reactions, representing possible mMoA, using the distance-based clustering and subnetwork extraction method. The first part of the workflow was exemplified on eight molecules selected for their known human hepatotoxic outcomes associated with specific MoAs well described in the literature and for which we retrieved primary human hepatocytes (PHH) transcriptomic data in Open TG-GATEs. Then, we further applied this strategy to more precisely model and visualize associated mMoA for two of these eight molecules (amiodarone and valproic acid). The approach proved to go beyond gene-based analysis by identifying mMoA when few genes are significantly differentially expressed (2 differentially expressed genes (DEGs) for amiodarone) or when very large number of genes were differentially expressed (5709 DEGs for valproic acid). In both cases, the results of our strategy well fitted evidence from the literature regarding known MoA. Beyond these confirmations, the workflow highlighted potential other unexplored mMoA.

**Conclusion:** The proposed strategy allows toxicology experts to decipher which part of cellular metabolism is expected to be affected by the exposition to a given chemical. The approach originality resides in the combination of different metabolic modelling approaches (constraint based and graph modelling). The application to two model molecules shows the strong potential of the approach for interpretation and visual mining of complex omics *in vitro* data. All code is freely available as well as data to reproduce results.

## 1 Introduction

Toxicology is entering a new era with the urgent need to follow a 3R (Reduce, Replace and Refine) policy when assessing risks of chemical molecules. The European Cosmetics regulation (EC) No 1223/2009 banning animal testing for cosmetic ingredients is a striking example of the need to develop non-animal approaches, particularly for systemic toxicity. In that context, new approach methodologies (NAMs) from the combination of *in silico* and *in vitro* methods are required to be fit for purpose and protective of human health. These NAMs are now being developed to support the so-called next generation risk assessment (NGRA) [1]. NAMs are already evolving at a fast pace thanks to the ever-increasing amount of omics data generated from *in vitro* experiments, creating unprecedented capacity to study biological systems. Omics screening allows, for instance, a more holistic classification of compounds based on their global effects [2,3] and it can be integrated to improve quantitative structure-activity relationships strategies [4]. The exploration of most biological processes is covered by transcriptomics or proteomics data. Among these processes, endogenous metabolism is becoming a source of concern since chemicals have the capacity to affect metabolism at a cellular and tissue level, potentially leading to adverse effects for humans such as diabetes, obesity, or even organ dysfunction [5–9]. Moreover, generating and analyzing several large-omics datasets can be expensive and methodologically challenging. Such datasets are usually of large dimensions (thousands of genes, proteins, and metabolites) that will contain information for broader insights on how an organism or a cell is globally impacted by a chemical. One of the current key challenges in the field is to extract knowledge from these rich but complex datasets.

The volumes of data generated by omics approaches and their complexity require advanced statistical and computational solutions to pinpoint patterns of interest among thousands of variables. Dimension reduction (PCA, MDS, t-SNE, etc.) methods are widely used to describe and visualize these large multidimensional datasets, but they are not designed for functional interpretation of omics data. From that perspective, enrichment-based methods, which allow the identification of biological modules (gene sets, metabolic pathways and cellular functions) in which identified modulated variables are over-represented, have the advantage of providing an overview of the studied biological modulations [10]. Nevertheless, these methods rely on arbitrary definitions of the pathways and functional sets, which might differ depending on the selected database, and tend to hide functional processes spanning several pathways [11,12]. Many studies [13–16] aim to take advantage of published gene expression data available in databases such as DrugMatrix [17], Connectivity Map [18,19], ToxicoDB [20] and Open TG-GATEs [21] (https://toxico.nibiohn.go.jp) to improve chemical toxicity assessment. For instance, Heusinkveld *et al.* [22] implemented an approach based on the comparison of Open TG-GATEs top 50 DEG signatures ranked according to their t-statistic. They aimed at providing both a score to compare compounds and a mechanistic understanding based on enrichment-based methods. On the other hand, Ting Li *et al.* [2] trained a deep neural network model for drug-induced liver injury prediction based on the LINCS L1000 dataset [19]. In both cases, metabolism is not the primary focus of the study, and the functional interpretation relies on enrichment-based methods, with the limitations explained above.

Therefore, to improve our understanding of how chemicals impact endogenous metabolism and can trigger potential adverse effects, it is necessary to develop new computational methods that would allow provision of functional metabolic information from omics data including the diversity of possible mMoAs leading to adverse outcomes. Genome-scale metabolic networks (GSMNs) are biological networks representing all the possible biochemical reactions occurring in an organism. They are therefore well suited to consider the diversity of endogenous metabolic disruptions potentially associated with adverse outcomes at a cellular or at an organ level. These networks are reconstructed from an annotated genome, curated, and refined thanks to biochemical knowledge retrieved from the literature [23,24]. Several GSMNs representing human metabolism such as Recon2 [25], Recon2.2 [26], Recon3D [27] and Human-GEM [28] have been published. GSMNs are composed of thousands of reactions and metabolites interconnected according to the stoichiometric matrix of the network. For instance, Recon2.2, which is one of the most used human GSMNs, is composed of 7785 reactions, 5324 metabolites, and 1675 associated genes distributed over 10 cellular compartments. The relationships between reactions and metabolites in the metabolic network are modelled by the stoichiometric matrix, which is the mathematical representation containing the proportions of substrates and products involved in each reaction. The stoichiometric matrix is the ground for constraint-based modeling approaches, which are largely used to model metabolic networks at the scale of cells, tissues, or organisms. Genes and reactions are linked by gene-protein-reaction (GPR) associations, which are formulated as Boolean rules, including all genes coding for either the isoenzymes (OR association) or the enzymatic complex subunits (AND association) catalyzing a reaction. Since these networks include all the reactions that can potentially be expressed in any tissue or cell type, and any condition for a given organism, they represent the global metabolism of this organism. It is then crucial to tailor these models to specifically represent the metabolism of one tissue, one cell, or one condition in order to accurately decipher the mMoA of chemical compounds.

Preliminary to any interpretation, it is essential to perform a condition-specific metabolic network reconstruction to avoid interpreting metabolic functions that would involve reactions that are not active in the biological condition or cell type under study. Condition-specific modeling approaches aim at exploiting experimental data to assemble a GSMN that more closely represents the condition under study. Many algorithms such as iMAT [29,30] or FASTCORE [31] have been developed to build condition-specific metabolic networks, most of which rely on transcriptomic data. Unlike other algorithms, these 2 algorithms do not require optimizing a physiological objective function such as biomass production maximization: this is especially more relevant for studying eukaryotic differentiated cells which are usually not growing anymore and might be oriented toward performing several other objective metabolic tasks.

FASTCORE aims to find a minimal flux consistent metabolic network that contains a list of “core” reactions and as few reactions from the global GSMN as possible. These “core” reactions correspond to all the reactions that are strongly assumed to be active in the studied condition and are defined by the user with the method of its choice, making FASTCORE a generic algorithm for reconstructing condition-specific metabolic networks [31]. The iMAT algorithm aims to find the best consensus between the reaction activity inferred from categorized gene expression data and the activity inferred from the GSMN structure, which defines the biochemical interdependencies between reactions through consumed and produced metabolites. The output is a condition-specific subnetwork that more faithfully represents the metabolic state of the cell in the condition of the transcriptomic experiment. Selecting the best algorithm among all these possibilities is not trivial as several benchmarking studies stated that none of the benchmarked methods outperforms the others for all cases [32] and that selection of the method should be mainly guided by the aim of the study and the available data [33].

One issue that is overlooked by most of these algorithms and that we tackled in our strategy is the alternative optimal solutions [34] issue. Indeed, due to the high complexity of GSMNs and the relatively low amount of biological data, the mathematical problem solved by algorithms such as iMAT and FASTCORE to find a condition-specific metabolic network is under-constrained. Practically, it means that many equally optimal condition-specific metabolic networks exist for a single biological condition and that arbitrarily taking one optimal condition-specific network would limit the reproducibility and bias further analyses made for that condition [34].

To avoid this problem, partial enumeration methods have been developed [34–37]. These methods explore the solution space (*i.e.*, the range of possible optimal condition-specific metabolic networks for this condition) to find a representative set of possible condition-specific metabolic networks. Rodriguez *et al.* formally described the enumeration problem by designing a network model with a finite and calculable set of alternative solutions which was used as a ground truth for evaluating partial enumeration performance. They highlighted that considering thousands of condition-specific metabolic networks is more robust and less error-prone than considering only one network in the solution space [34]. Nevertheless, it complexifies the interpretation since a condition will be associated with thousands of potential network configurations. There is therefore a need to define a new strategy to extract mechanistic information from these numerous condition-specific metabolic networks to allow the final identification of the mMoA.

In this study, we propose a strategy designed to reduce the increased analysis complexity resulting from constraint-based modelling coupled with partial enumeration by calculating DARs and exploiting them with graph analysis methods with the final aim to predict and visualize chemicals’ mMoA. To develop our strategy, we used gene expression data from the Open TG-GATEs database and we built condition-specific metabolic networks with a partial enumeration method adapted from the DEXOM [34] approach. Open TG-GATEs has been created from two projects spanning several years: Toxicogenomics Project One [38] (TGP1: 2004 to 2007) and Toxicogenomics Project Two [39] (TGP2: 2010 to 2011). Open TG-GATEs contains gene expression data generated on Crl:CD Sprague–Dawley rats and on primary human hepatocytes (PHH) and primary rat hepatocytes, after exposure to low cytotoxic doses of 150 pharmaceutical molecules. We selected gene expression data generated on PHH exposed for 24h to the maximum dose, which corresponds to a dose inducing less than 20% of cytotoxicity. In order to evaluate the metabolic impact of a given chemical at a specific dose, we developed a statistical approach consisting of identifying DARs between two distinct sets of condition-specific metabolic networks representing respectively the exposure condition and the control (*i.e.,* without exposure) condition. We took advantage of the GSMN structure to compute the metabolic distance between DARs in order to identify clusters of functionally interdependent metabolic reactions. We then extracted a minimal subnetwork for each cluster to better understand how these small biological processes are constructed and how reactions are interconnected. Finally, we used MetExploreViz [40] to interactively visualize the subnetworks and overlay additional annotation on the network such as cellular compartments, metabolic pathways, and custom mappings. This approach enables the analysis of the metabolic impact of a chemical at a global level but also at a very precise level such as the biochemical reaction scale. The workflow has been applied to eight molecules (ethanol, valproic acid, indomethacin, amiodarone, allopurinol, rifampicin, sulindac, and tetracycline) selected for their known hepatotoxic effects and the large amount of scientific literature describing their MoA. We further highlight capabilities of our strategy by predicting and analyzing the metabolic impact of two known hepatotoxic compounds: amiodarone and valproic acid.

## 2 Results

We set up a strategy to better understand the mMoA of chemicals through transcriptomic data integration with condition-specific modelling enhanced by partial enumeration: this strategy combines constraint-based modelling methods and graph technique in a three-steps workflow.

The first step (Fig 1, box A) consists of building condition-specific metabolic networks. Indeed, GSMNs encompass all possible metabolic reactions regardless of the tissue, cell, or condition. Performing computational analysis on this generic model may raise inaccurate results (*e.g.*, highlighting a pathway which is known to be inactive for a cell type). Condition-specific metabolic networks are composed of reactions predicted as active by a modeling algorithm for a given biological condition based on the generic GSMN and a set of gene expression data. In this study, we used Recon2.2 [26] as the initial GSMN to build reconstructions for several biological conditions (*i.e.*, cells exposed to a chemical) and transcriptomic data obtained from the Open TG-GATEs database (described in “Methods” in the section “Transcriptomic data processing”). For each studied condition, a set of condition-specific metabolic networks optimally matching gene expression data, network topology and the stoichiometry of reactions is computed with an adapted version of DEXOM [34]. Our major adaptation of the DEXOM method consisted in combining 2 of the proposed algorithms (reaction-enum followed by diversity-enum) while reducing the number of enumerated solutions by randomly selecting starting solutions for diversity-enum among all the reaction-enum solutions, rather than using all of them (details are provided in the “Methods” section “). Although our adapted DEXOM version provides a more partial enumeration, it allows to reduce the computational cost. The output is a dataset that gathers, for each reaction, the number of times it has been predicted as active by the algorithm across all optimal enumerated solutions. This result highlights the likelihood of each reaction of being active for a specific condition. Note that this step allows the prediction of activity for reactions, even if they are not associated with any gene (*e.g.*, passive transport reactions), by inferring their activity from the activity of surrounding reactions.

**Fig 1.**
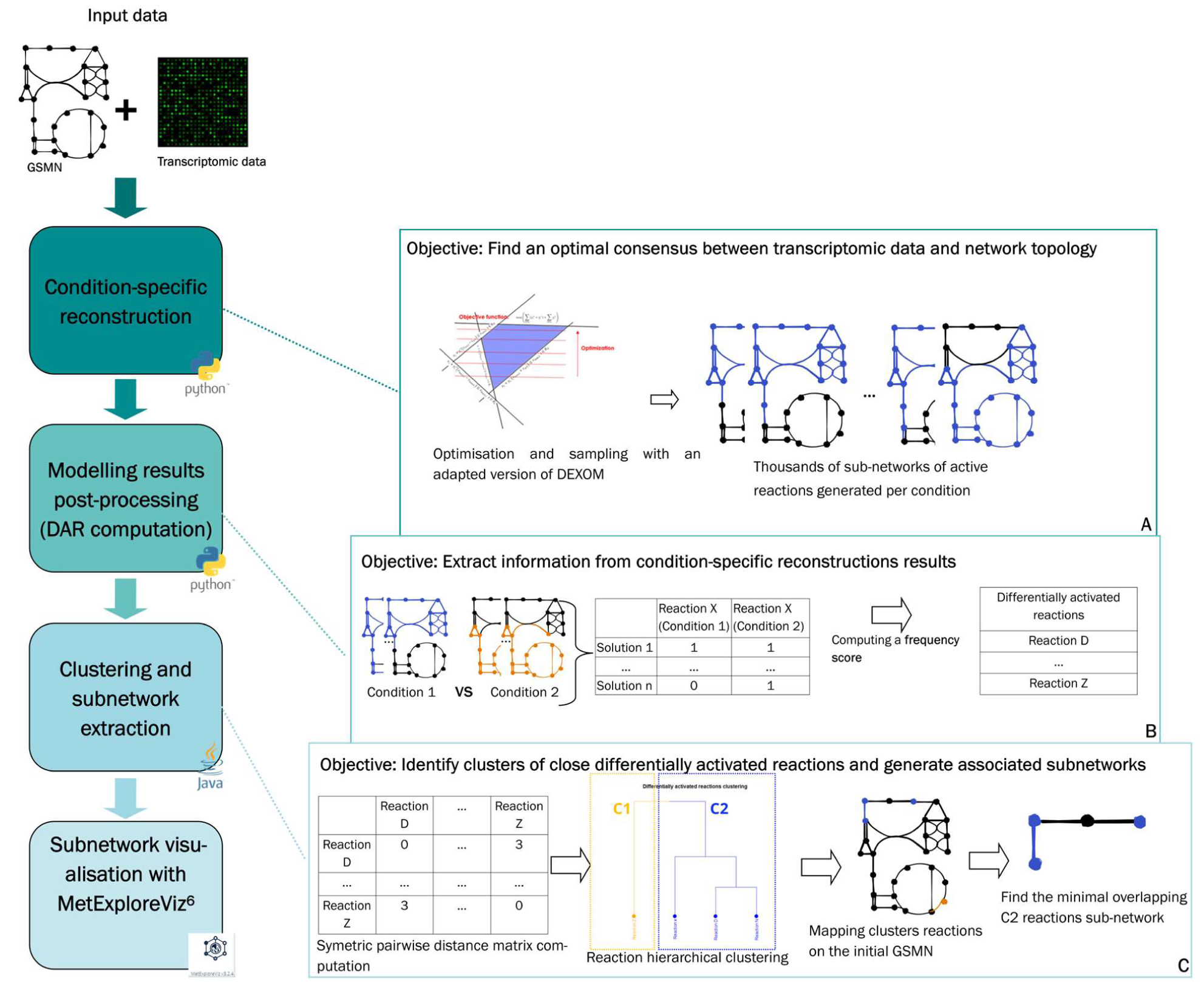
General overview of the three-step strategy. In the first step (A), transcriptomics data are integrated to a GSMN with a partial enumeration approach adapted from DEXOM. In the second step (B), DARs are computed from the large number of metabolic networks obtained after the optimization and sampling step. Finally, in the third step (C), network analysis methods are developed and employed to interpret DARs and improve our understanding of the chemicals’ mMoA.

In the second step (Fig 1, box B) of the workflow, the objective is to identify reactions which activity changes in a significant manner between control and treated conditions. We therefore apply a statistical test between sets of reactions’ predicted activity in order to identify DARs between the two conditions (described in “Methods” in the section “Identification of differentially activated reactions”). Since some reactions cannot be correctly constrained by transcriptomic data (*e.g.*, associated with less specific genes or no gene at all), they are more likely to be indiscriminately predicted as active or inactive while maintaining the same optimal fit with the data. The predicted activity of these reactions may therefore be more variable even across optimal subnetworks representing the same condition. We implemented a “baseline noise” calculation and filtration approach (described in “Methods” in the section “Baseline noise calculation”) to remove these reactions from the list of DARs obtained at the end of this step and to ensure their robustness. The list of DARs by itself is a very insightful result since it allows the listing of reactions whose activity is significantly affected by the studied condition (exposed *vs* control cells).

The last step (Fig 1, box C) of the workflow consists of deciphering where and how cell metabolism is perturbed by chemical exposure. DARs are not independent and may act in a coordinated manner through sequences of reactions, hence highlighting the potential modulated cascade of enzymatic reactions. To do so, a network-based approach has been developed to detect, visualize, and analyze the functional role of previously identified DARs. The method aims at stratifying the list of DARs into clusters of close reactions in the metabolic network (detailed in “Methods” in the section “DARs clustering”). Two reactions are considered to be close when they are connected to each other with only a few intermediary reactions. Once these clusters were identified, close reactions were used to extract small human-readable subnetworks (detailed in “Methods” in the section “Subnetwork extraction”) that would describe parts of the metabolic network that are specifically modulated in the studied condition, suggesting potential mMoA of the molecules.

### Identification of differentially activated reactions (DAR) associated with exposure to eight chemicals

The strategy has been applied to eight molecules present in the Open TG-Gates database (ethanol, valproic acid, indomethacin, amiodarone, allopurinol, rifampicin, sulindac, and tetracycline) selected for their known liver toxicity and the associated MoAs reported in the literature. We used gene expression data generated in PHH exposed for 24h at the highest concentration of each of these 8 molecules along with their associated controls. Condition-specific metabolic networks were reconstructed (Fig 1, box A) from gene expression data for each retrieved sample (two replicates per condition). On average, 10,000 alternative optimal solutions were enumerated for each sample. Each optimal solution is composed of a unique set of active and inactive reactions. In Table 1, the minimal, maximal, and average number of active reactions for the solutions calculated by DEXOM for each chemical and two vehicle controls (*i.e.*, medium and DMSO) are reported. It is worth noting that the size (*i.e.*, the number of active reactions) of DEXOM optimal solutions is in the same order of magnitude for all conditions (treatment or controls, see Table 1) with a minimal number of active reactions ranging from 3423 (DMSO condition) to 3639 (tetracycline 25µM, 24h), a maximal number of active reactions ranging from 4396 (valproic acid 5000µM, 24h) to 4645 (indomethacin 200µM, 24h), and a mean number of active reactions ranging from 4070 (valproic acid 5000µM, 24h) to 4570 (amiodarone 7µM, 24h).

**Table 1.**
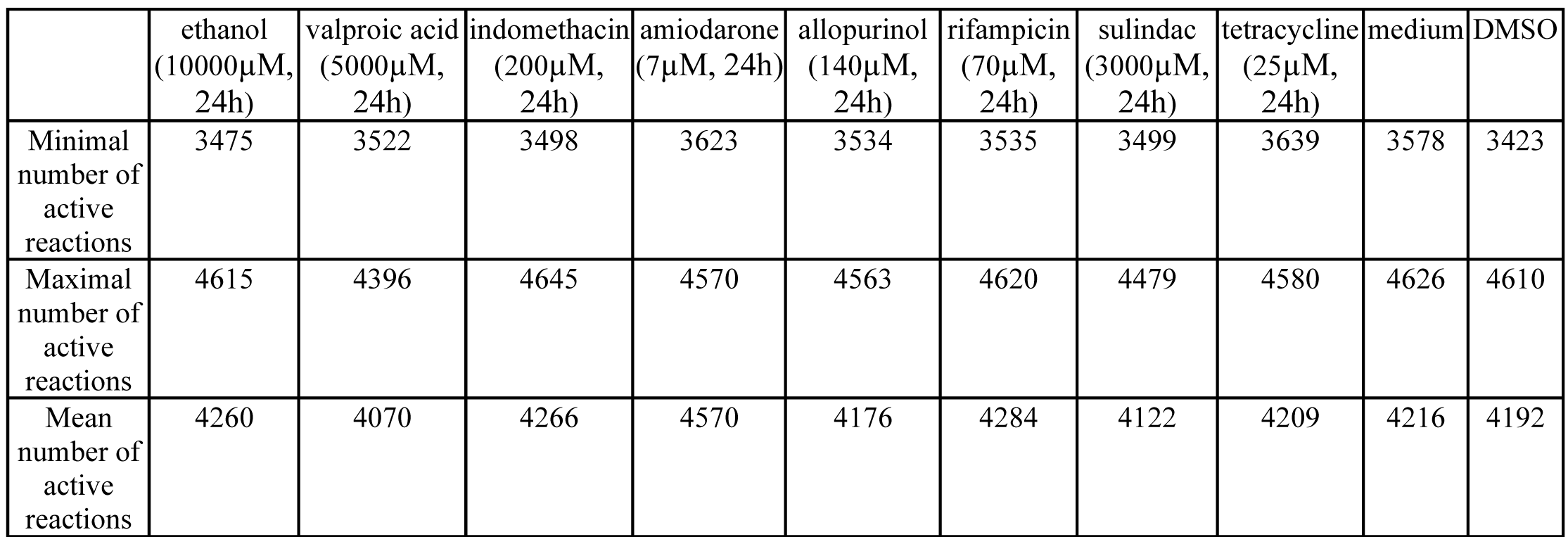
Number of predicted active reactions for each chemical and vehicle set of condition-specific metabolic networks. Sets of condition-specific metabolic networks were calculated by the adapted DEXOM approach implemented in our workflow (Fig 1, box A). All solutions obtained for samples of the same molecule have been merged before calculating the minimal, maximal, and mean number of active reactions per condition.

DARs were computed to identify metabolic reactions modulated following exposure to each selected molecule (Fig 1, box B) compared to the corresponding control condition. The number of identified DARs ranged from 57 for PHH exposed to indomethacin (200µM, 24h) to a maximum of 477 for PHH exposed to valproic acid (5000µM, 24h) (Table 2). Valproic acid, indomethacin, amiodarone and allopurinol DARs lists were only slightly affected by the noise filtration procedure with 12.6%, 5.3%, 6.7% and 11.4%, respectively, of DARs filtered out (Table 2). Conversely, for tetracycline and ethanol, 43.4% and 62.8% of DARs were respectively filtered out (*i.e.*, reactions being modulated by both molecules and control conditions (S1 Table)). For each condition, we also computed the DARs specificity ratio as the number of DARs retrieved solely in this condition and not in any other studied condition (*i.e*., DARs specific to the condition) divided by the total number of DARs retrieved for this condition (S2 Table). Calculated DARs specificity ratios ranged from 22.2% for indomethacin to 91.1% for valproic acid, indicating that the predicted mMoA for indomethacin was at least similar to one other condition in the study, whereas in contrast, the predicted mMoA for valproic acid was quite different from any other condition in the study.

**Table 2.** Number of DARs obtained for each condition. DARs were identified by the developed workflow for PHH after 24h exposure to eight molecules known for their hepatotoxicity at dose levels yielding an 80-90% survival ratio.

| | ethanol<br>(10000μM,<br>24h) | valproic acid<br>(5000μM,<br>24h) | indomethacin<br>(200μM,<br>24h) | amiodarone<br>(7μM, 24h) | allopurinol<br>(140μM,<br>24h) | rifampicin<br>(70μM, 24h) | sulindac<br>(3000μM,<br>24h) | tetracycline<br>(25μM, 24h) |
| --- | --- | --- | --- | --- | --- | --- | --- | --- |
| Number of DARs | 94 | 477 | 57 | 60 | 88 | 121 | 242 | 99 |
| Number of DARs after filtration | 35 | 417 | 54 | 56 | 78 | 98 | 181 | 56 |
| % of reactions filtered out | 62.8 | 12.6 | 5.3 | 6.7 | 11.4 | 19 | 25.2 | 43.4 |

### Global analysis of DARs associated with exposure to amiodarone and valproic acid

Two of the eight molecules were selected for further analysis: amiodarone (7µM, 24h) and valproic acid (5000 µM, 24h). These two molecules were selected because 1) they are well described hepatotoxic compounds with several MoAs known to induce liver damage such as steatosis [41,42], and 2) their level of transcriptomics-based information differs, with 5709 and two differentially expressed genes (DEGs) for PHH exposed to valproic acid and amiodarone, respectively (described in “Methods” in the section “DEG identification”). Despite the low number of DEGs observed for amiodarone, 56 DARs were identified in PHH after 24h exposure to 7µM amiodarone using the developed method.

Fig 2 shows the visualization produced by MetExploreViz for DARs identified for amiodarone and valproic acid in the context of human GSMNs using Recon 2.2. For amiodarone (Fig 2 A), a main group is composed of well interconnected up-activated reactions, while few other groups of up- or down-activated DARs are disconnected with no shared substrates or products. For valproic acid (Fig 2 B), many small groups of DARs are disconnected from each other without any particular area of the metabolic network specifically impacted. Hence, this general observation indicates that the mMoAs might differ between these two molecules, and further in-depth analysis is required.

**Fig 2.**
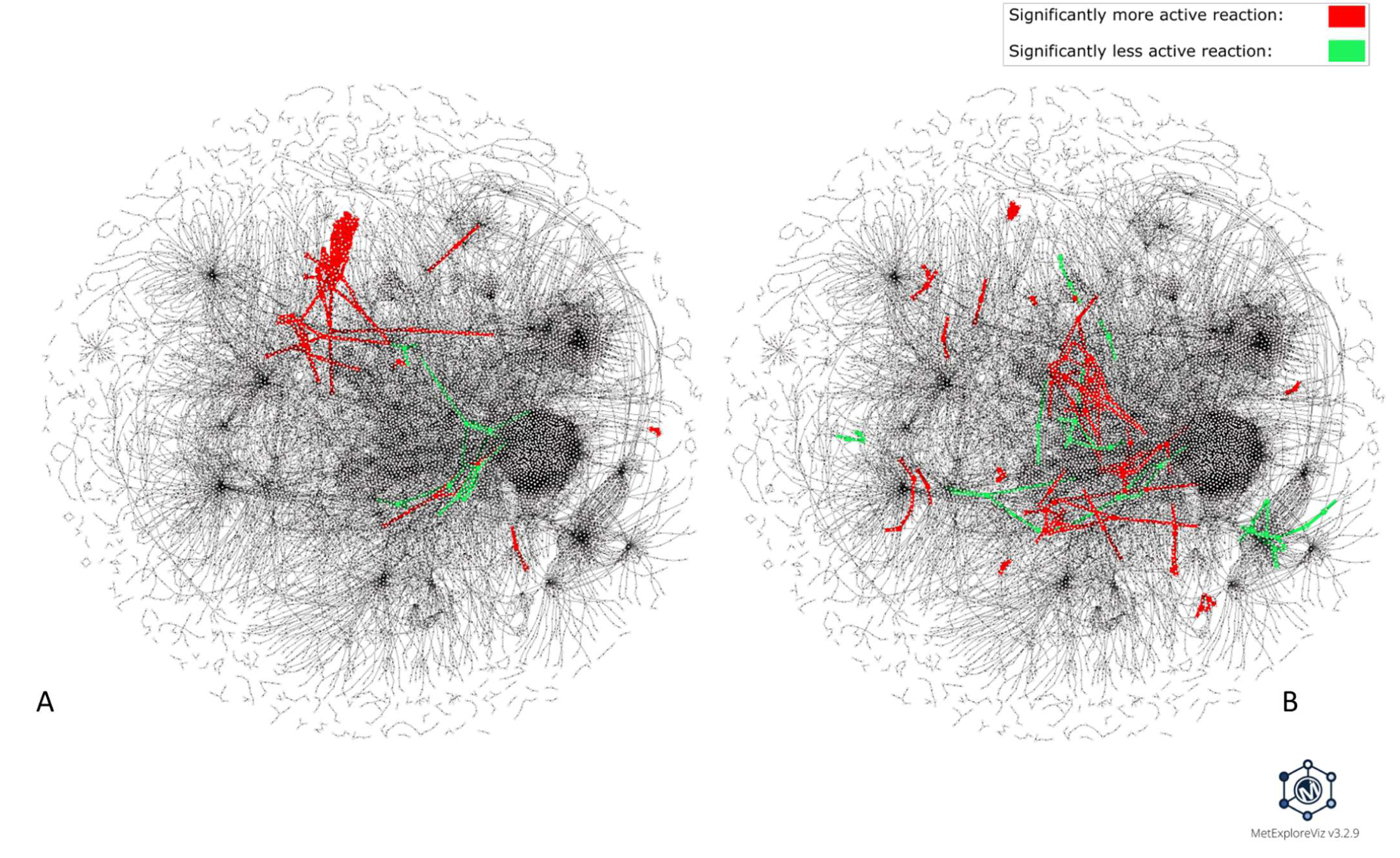
Visualization of DARs identified for amiodarone and valproic acid within the Recon2.2 metabolic network. This visualization was performed with MetExploreViz while removing side compounds (S3 Table). DARs identified for amiodarone (7µM, 24h) are highlighted in Fig 2 A and DARs identified for valproic acid (5000µM, 24h) are highlighted in Fig 2 B, with more frequently active reactions in the exposed *vs* control condition colored in red and less frequently active reactions in green. Nodes represent reactions and metabolites and are connected if a metabolite is a product or a substrate of a reaction. Fig 2 A and Fig 2 B are both based on the same network layout; thus, each reaction and metabolite is located at the same coordinates, allowing visual comparison.

### Identifying subnetworks of DARs with graph-distance clustering and subnetwork extraction

In order to capture the mMoA of a molecule without having to rely on subjectively defined pathways [11,12], it is assumed that the closeness within the network can be used as a measure of the metabolic proximity between reactions. Indeed, reactions are linked through compounds being produced and consumed by others. The shorter the chain between two reactions, the stronger the expected interdependency. Therefore, the metabolic distance, which is the length of the chain or path between two reactions in the metabolic network, is used to estimate interdependency. Hence, we propose grouping reactions that are closely located as a proxy to identify reactions involved in the same metabolic function.

The pairwise distance matrix between all reactions identified as differentially active for each of the two molecules, amiodarone and valproic acid, was computed based on the list of DARs. As described in the “Methods” section, the computation of pairwise distances was performed on the metabolic reaction undirected graph of Recon2.2, using the shortest path distance metric. Then, subsets of DARs were identified by clustering the pairwise distance matrix with a hierarchical clustering approach. Two clusters were selected for PHH exposed to 7µM amiodarone for 24h (Fig 3 A) and three clusters for PHH exposed to 5000µM valproic acid for 24h (Fig 3 B).

**Fig 3.**
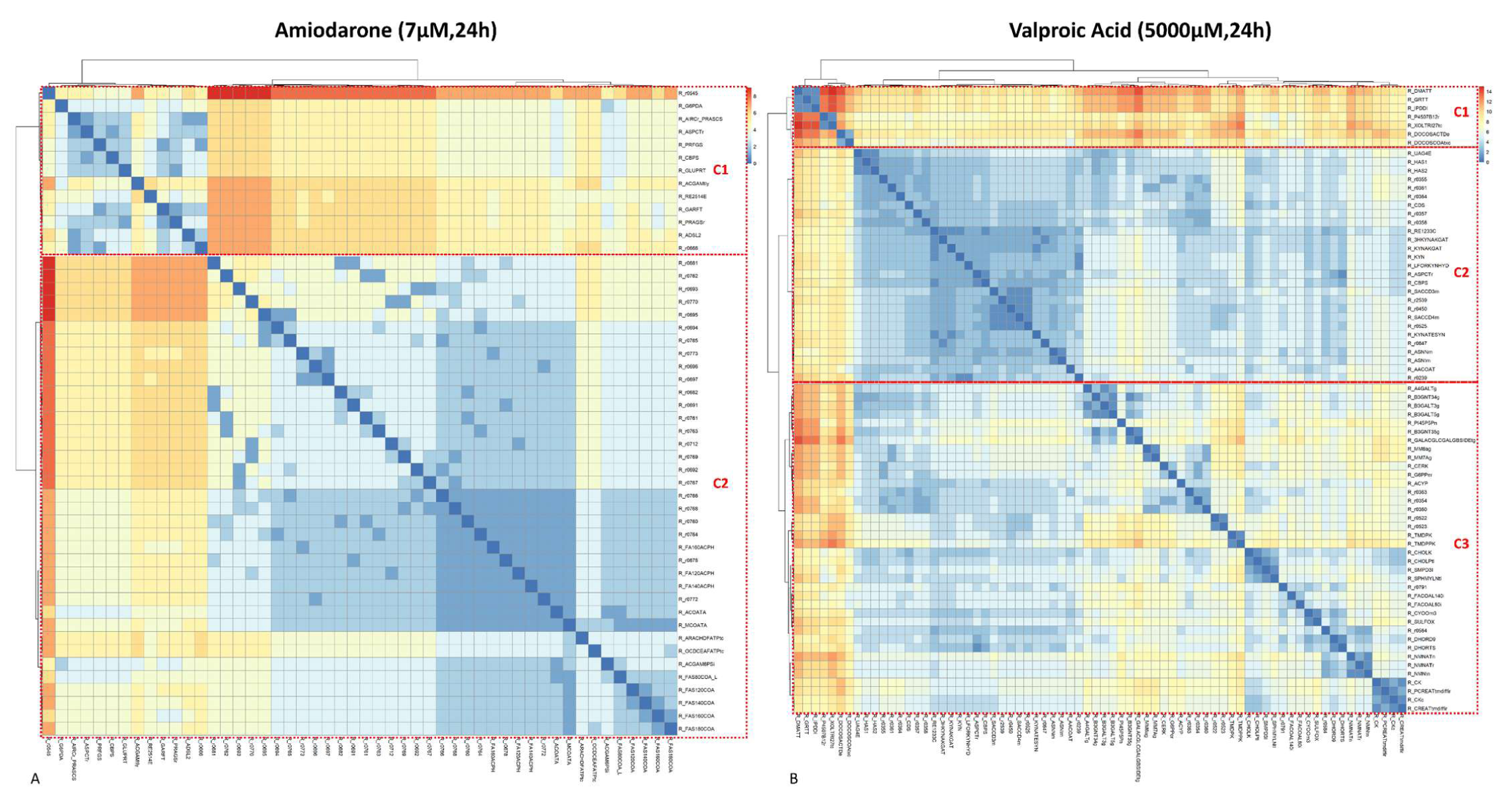
Biclustered heatmap of the pairwise reaction distance matrix for amiodarone and valproic acid. Hierarchical biclustering on the pairwise reaction distance matrix for amiodarone and valproic acid was performed with the Ward algorithm. The resulting biclustering was visualized as a heatmap computed with the Pheatmap R package. The color scale depicts the distance between two reactions. The distance ranges between zero (cells colored in blue) to eight for amiodarone (Fig 3 A) and 14 for valproic acid (Fig 3 B) (cells colored in red). Two main clusters (C1 and C2) were identified for amiodarone (Fig 3 A) and three clusters (C1, C2, and C3) were identified for valproic acid (Fig 3 B).

To go further in the interpretation and to better understand how these reactions interact and are implicated in the mMoA of our chemicals of interest, a subnetwork extraction step was implemented and applied to each cluster of DARs. Extracting a subnetwork allows one to visualize how the DARs are interconnected and fill gaps between disconnected DARs by adding the necessary intermediate reactions. The subnetwork extraction was performed using the Met4J library (https://forgemia.inra.fr/metexplore/met4j) implementation of the minimal Steiner tree on a pruned reaction graph of Recon2.2. The minimal Steiner tree extraction algorithm enables the extraction of a subnetwork connecting two lists of nodes (*i.e.*, DARs) while minimizing the size of the final subnetwork. Therefore, this algorithm well fits our objective of visualizing interactions between DARs while keeping it human-readable. To guide our analysis, the “DARs subnetwork coverage” metric was defined, which is calculated as the number of DARs in a subnetwork divided by the total number of reactions in this subnetwork. This metric provides an estimate of how rich in DARs a subnetwork is. A high DARs subnetwork coverage score indicates closely located and tightly interacting DARs.

For PHH exposed to 5000µM of valproic acid for 24 hours, a subnetwork with a 77% DARs subnetwork coverage was extracted (Cluster C2 in Fig 3B, see S4 Table for details), meaning that most reactions within the extracted subnetwork were DARs that were closely interconnected. Among the DARs of this subnetwork, 21 were up-activated while only four were down-activated (Fig 4A). Five DARs predicted as up-activated were associated with -oses metabolism such as fructose/mannose metabolism and glycolysis/gluconeogenesis pathway, some of which are involved in the phosphorylation of hexoses (Fig 4B). Another group of up-activated DARs is associated with lysine metabolism in mitochondria (html links for visualization in S1 Appendix) and especially the degradation of lysine through production of L-saccharopinate from 2-oxoglutarate. One up-activated reaction is associated with transport of L-asparagine into the mitochondria, which is then used by the L-asparaginase to produce L-aspartate. The resulting L-aspartate is finally transported by an aspartate-glutamate shuttle that was not differentially modulated. The phosphatidate cytidylyltransferase associated with the glycerophospholipid metabolism was also predicted to be up-activated in PHH after exposure to 5000 µM valproic acid for 24 hours. Finally, eight differentially up-activated reactions were associated with tryptophan metabolism and involved in reactions associated with kynurenate, L-glutamate and 2-oxoglutarate. Down-activated reactions were more sparsely located in the extracted subnetwork and were associated with hyaluronan metabolism, propanoate metabolism, where acetoacetate and coenzyme A are conjugated to produce acetoacetyl-CoA, and finally with pyrimidine synthesis, where glutamate and aspartate production/consumption were disturbed. The subnetwork extraction algorithm could add reactions that were not identified as DARs but were necessary for the connectivity of the subnetwork. These added reactions were associated with pathways also associated with DARs such as mitochondrial transport, glycolysis/gluconeogenesis, pyrimidine synthesis and lysine metabolism (Fig 4B).

**Fig 4.**
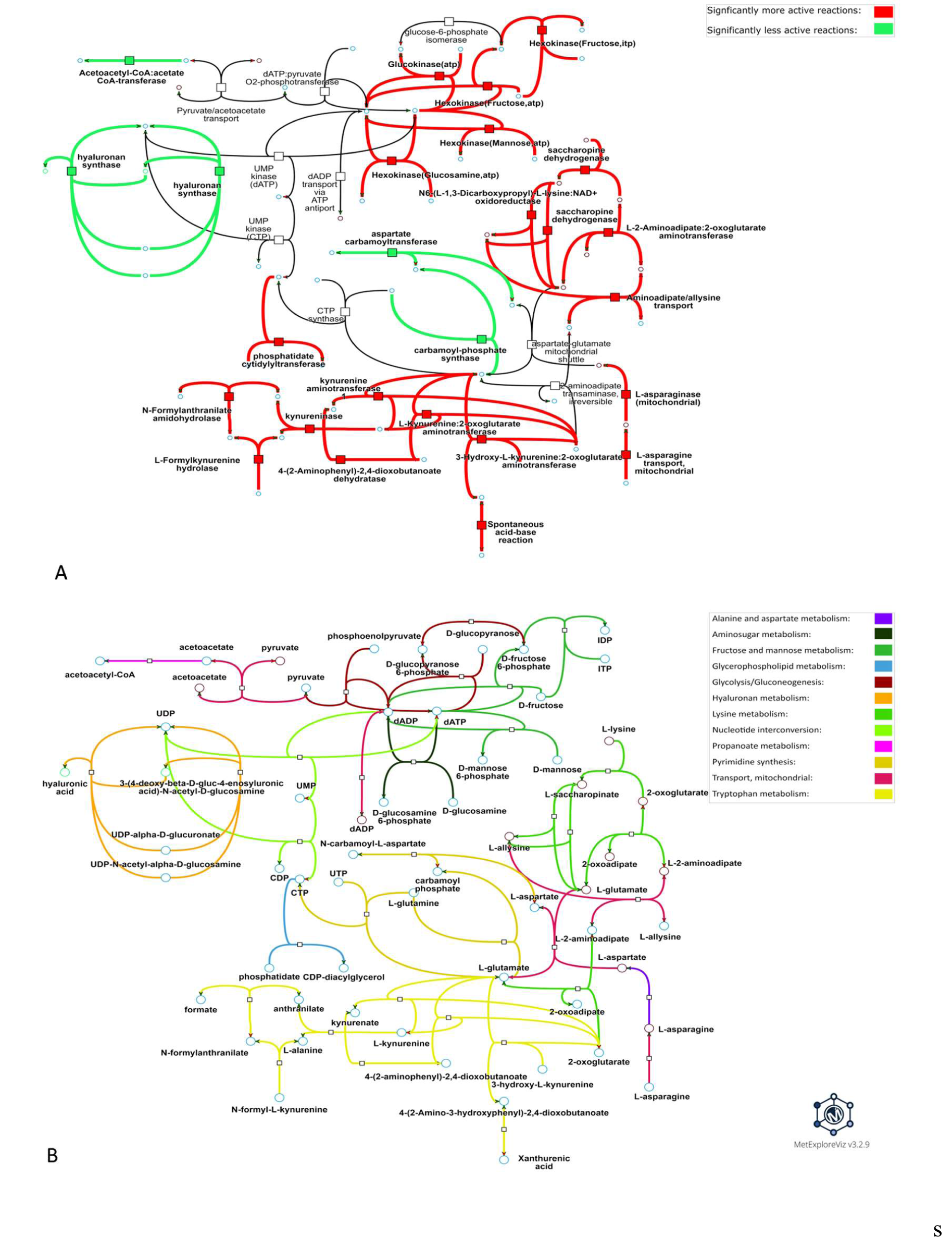
Metabolic visualization of the metabolic subnetwork extracted from one of the DARs clusters predicted for valproic acid. DARs were predicted by performing condition-specific reconstructions for PHH exposed to 5000µM valproic acid for 24h. The visualized subnetwork was computed from DARs in cluster #2, which is the cluster with the highest DARs subnetwork coverage among the clusters identified in the distance matrix (see Fig 3 B) for this condition. Nodes represented by a square are metabolic reactions and nodes represented by a circle are metabolites. Fig 4A and 4B represent the same subnetwork with the same topology. Links in Fig 4A are highlighted according to the direction of the perturbation (*e.g.*, if the reaction is more frequently active in the exposed *vs* control condition) and links in Fig 4B are colored according to the metabolic pathway of the associated reaction. Interactive visualizations can be accessed through the following links: https://metexplore.toulouse.inrae.fr/userFiles/metExploreViz/index.html?dir=/72ff7fdc7031b880ef4f3532134aa326/networkSaved_292937465 and https://metexplore.toulouse.inrae.fr/userFiles/metExploreViz/index.html?dir=/72ff7fdc7031b880ef4f3532134aa326/networkSaved_1994092833

For PHH exposed to 7µM of amiodarone for 24 hours, a subnetwork with a DARs coverage of 95% (S4 Table) was extracted, indicating that nearly all the reactions that make up this subnetwork are DARs. 36 DARs is up-activated in the treated condition and only one DAR is down-activated in the treated condition (Fig 5A). Most of the up-activated reactions (*i.e.*, 31) of this subnetwork are associated with the fatty acid synthesis, four reactions are linked to the fatty acid oxidation pathway and one reaction is associated with aminosugar metabolism. Regarding the fatty acid synthesis pathway, many reactions associated with conjugation/deconjugation of acyl carrier protein (ACP) to fatty acids were perturbed. Fatty-acyl-CoA synthase reactions were also perturbed with many reactions involved in the addition of malonyl-CoA to fatty acids as being differentially up-activated. Some up-activated reactions associated with the fatty-acid oxidation pathway were also disturbed, such as the acetyl-CoA-ACP transacylase and the malonyl-CoA-ACP transacylase as well as two fatty-acyl-ACP hydrolases. Finally, the only down-activated reaction of this subnetwork is a reaction associated with the aminosugar metabolism involved in N-acetylglucosamine-6-phosphate’s production from acetyl-CoA and D-glucosamine-6-phosphate. In the subnetwork presented in Fig 5, only two reactions associated with fatty-acyl-CoA elongation from the fatty-acid synthesis pathway were added by the subnetwork extraction algorithm.

**Fig 5.**
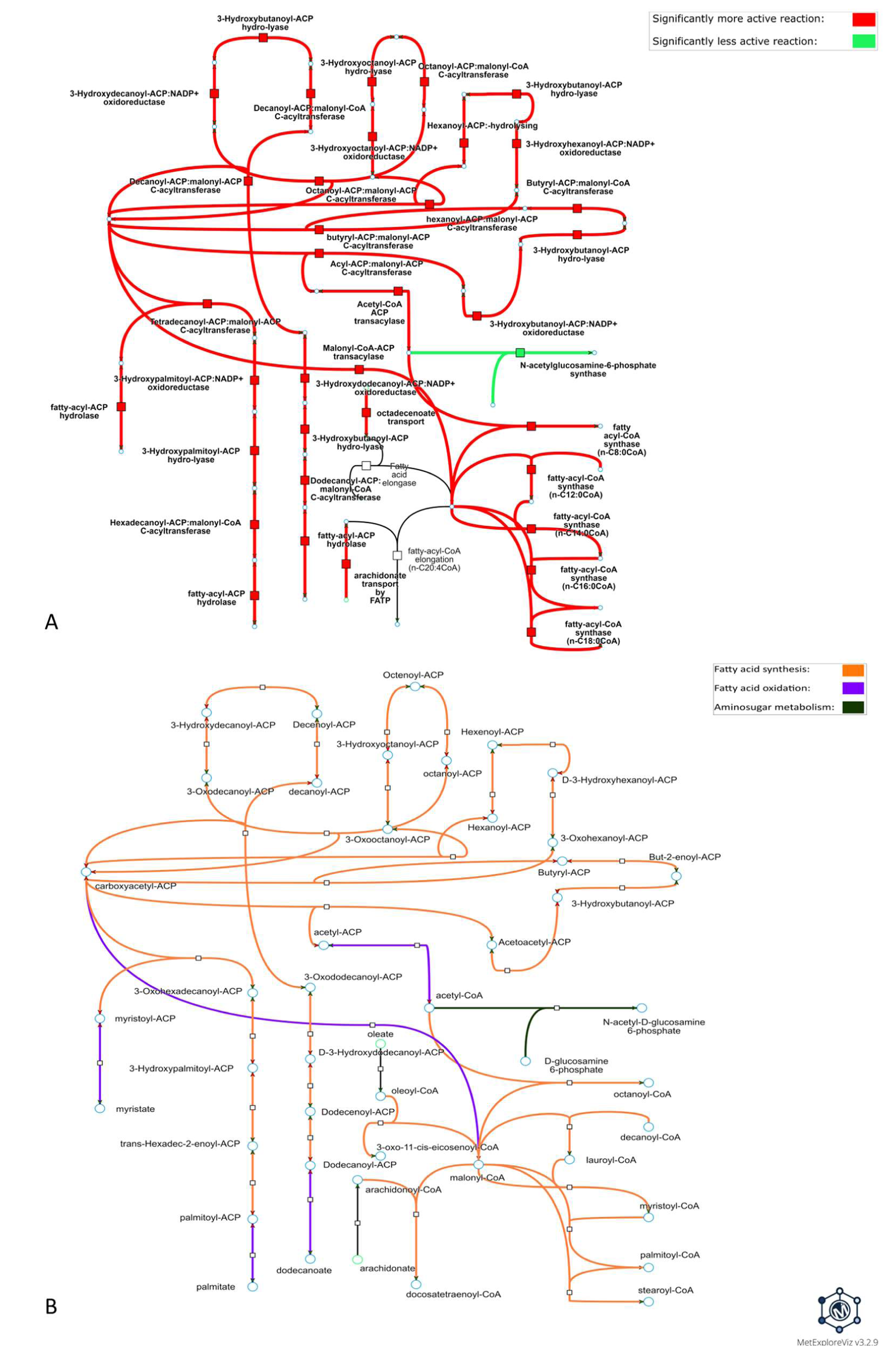
Metabolic visualization of the subnetwork extracted from one the DARs cluster predicted for amiodarone. DARs were predicted by performing condition-specific reconstructions for PHH exposed to 7µM amiodarone for 24h. The visualized subnetwork was computed with DARs from cluster #2, which is the cluster with the highest DARs subnetwork coverage among the clusters identified in the distance matrix (Fig 3A) for this condition. Nodes represented by a square are metabolic reactions and nodes represented by a circle are metabolites. Fig 5A and 5B represent the same subnetwork with the same topology. Links in Fig 5A are highlighted according to the direction of the perturbation (*e.g.*, if the reaction is more frequently active in the exposed condition *vs* control condition) and links in Fig 5B are colored according to the metabolic pathway of the associated reaction. Interactive visualizations can be accessed through these links: https://metexplore.toulouse.inrae.fr/userFiles/metExploreViz/index.html?dir=/72ff7fdc7031b880ef4f3532134aa326/networkSaved_373423088 and https://metexplore.toulouse.inrae.fr/userFiles/metExploreViz/index.html?dir=/72ff7fdc7031b880ef4f3532134aa326/networkSaved_725935955

## 3 Discussion

### Choosing a modelling algorithm able to explore alternative solutions

The proposed strategy has been applied to Recon2.2 for demonstration purposes. It can also be used with other Human GSMN like HumanGEM, Recon3D or any another GSMN following SBML standard. In fact, the only requirement is to be able to map transcriptomics data using adequate identifiers for gene expression integration. Choosing which GSMN to use among the variety of published GSMNs can be a challenging task due to the difficulty to compare these metabolic networks [43,44]. To our knowledge, no standard procedure exist to compare in an objective manner GSMNs [45]. According to [43,44], the main challenge to compare GSMNs is the lack of a standardized procedure, the lack of published differences between GSMNs, and the lack of common identifiers. Therefore, each author should define their own criteria for selecting the best GSMN to answer the question raised in their study.

Integration of gene expression data into GSMNs allows additional information to be uncovered that could not be retrieved from transcriptomics data, such as the activity of reactions not associated with the expression of specific genes (passive transports, spontaneous reactions, etc.) and cellular localization. To be predicted as active, a reaction must have its source metabolite(s) being produced by another reaction in the metabolic network, and its product(s) being consumed by another reaction. Therefore, the objective of the modeling algorithm is to find the best consensus between reactions predicted as active according to gene expression data and these production/consumption constraints from the metabolic network.

As mentioned previously, the amount of information available in gene expression data for the modeling is not sufficient to allow the algorithm to find a unique condition-specific network. This limit has already been described by several authors [34–36,46] and Rodriguez *et al.* demonstrated that the analysis of a single solution as it is usually done with standard implementations of widely used constraint-based algorithms (*i.e.* iMAT, FASTCORE, GIMME,…) return less robust results [34]. To circumvent this issue, partial enumeration approaches have been developed [34,35,37,46] to obtain a set of alternative solutions that is representative of the diversity of solutions existing in the solution space. Such methods provide many similarly adequate optimal subnetworks based on the gene expression data. Rossell *et al.* proposed EXAMO to reconstruct a minimal condition-specific network from the alternative optimal solution space that prioritizes reactions found to be active in all alternative solutions. This strategy tends, however, to emphasize central metabolism although it might not necessarily be the part of metabolism most affected by chemical compound exposure. Poupin *et al.* identified required, inactive, and potentially active reactions, based on how frequently the reactions were found to be active in the alternative optimal solution space. In this method, the group of reactions defined as potentially active is difficult to interpret and ended up being merged with required reactions to perform the pathway enrichments analyses in their study. MOOMIN [37] also provides an alternative approach able to explore the solution space, by predicting whether reactions are likely to be up or down activated based on differential expression data.

In our strategy we chose to use the DEXOM method for its ability to explore the solution space and to return a set of alternative solutions that were shown to represent the diversity of solutions that can be found in the solution space [34]. DEXOM also has the advantage to output a qualitative prediction of reactions activity, not relying on computed metabolic flux values. Indeed, several studies [47–49] reported a low correlation between gene expression intensities and metabolic fluxes values, whereas other studies [32,50] showed that gene expression data can be used as a good proxy for predicting metabolic fluxes. Given the lack of consensus and formal proof on a direct and quantitative relation between gene expression data and metabolic fluxes, the strategy adopted in DEXOM consists in binarizing flux values and considering a reaction as active if it is predicted to carry a non-zero flux.

As discussed above, exploring the alternative solution space is important to obtain more robust results. A complementary approach not explored in this study is to further constrain the space of solutions by combining gene expression data with medium metabolomics data (exometabolomics) that would add further constraints on metabolic fluxes, thereby reducing the uncertainty related to the number of possible alternative solutions and obtain more accurate metabolic fluxes predictions.

### Identification of DARs from the partial enumeration of condition-specific reconstructions

Using DEXOM to explore the alternative solution space of each modelled condition is crucial to increase the robustness of the results. However, the subsequent results analysis is a challenging task because, for each condition, thousands of different condition-specific metabolic networks need to be analyzed. To solve this challenge, we introduce the DAR calculation approach, which aims to compare two sets of condition-specific metabolic networks and identify reactions that are significantly perturbed in one condition compared to the other one (*e.g.*, treated *vs* control). However, some metabolic reactions are more sensitive to the under-constrained problem and will be predicted as active or inactive independently of experimental data specificity. To limit this effect, we added a filtration step called “baseline noise filter” that takes into account the uncertainty of predicting the activation status associated with some reactions. The baseline noise filter had a different impact on DARs lists depending on the condition. PHH exposed to ethanol was the most impacted condition with 62.8% of its DARs removed by the filter, while PHH exposed to indomethacin was the least impacted condition with 5.3% of its DARs removed. Most of the DARs filtered out for ethanol and tetracycline (*i.e.*, the most impacted conditions) are transport and peroxisomal β- oxidation reactions. Indeed, these reactions are more prone to noisy predictions either because they are not associated with any genes (*e.g.*, exchange reactions) or they are associated with complex GPR rules that tend to result in unconstrained reactions. The baseline noise filter is a rather conservative approach, but it ensures that the predicted

perturbation is not due to a lack of constraints on the model. Indeed, one key point in the development of condition-specific reconstructions was to get enough biological constraints. Both the DAR computation and the baseline noise filtration steps implemented in our workflow can be adapted to other partial enumeration approaches as they only require activation frequencies for each metabolic reaction in two distinct conditions.

### Comparison and complementarity with pathway enrichment

To evaluate the added value of our workflow, in comparison with directly using gene expression data through approaches such as pathway over-representation on DEG, we performed a pathway over-representation analysis (ORA) on DEGs for the valproic acid condition (S1 Fig). The first limit of this DEG-based approach is that it becomes inapplicable or unreliable when the list of DEGs is short, which is often the case for cells exposed to a low concentration of chemicals triggering subtle metabolic impacts. It is indeed the case in our study for the 24h treatment of PHH with 7µM amiodarone, where only two DEGs were evidenced. On the contrary, we showed that using our strategy, it is possible to analyze the metabolic impact of a chemical starting from transcriptomic data even when the DEGs list is short. Another limit of using DEG-based approaches to explore metabolic alterations is that it provides information on changes that are often not directly link to metabolism but involve more generic cellular processes. For instance, when performing pathway ORA on DEGs with Reactome (*i.e.*, including all genes, not just metabolic genes) for PHH exposed to 5000µM valproic acid for 24h, we observed that many of the top 50 enriched terms are related to cell cycle, gene expression regulation, and cell signaling (S1 Fig). Although this is valuable information about the MoA of valproic acid, it is not directly associated with its metabolic impact in liver cells, suggesting that combining both gene expression data analysis and condition-specific reconstruction is beneficial. We also performed a pathway ORA with DARs on Recon2.2 pathways (S2 Fig) for the eight molecules. For amiodarone, predicted DARs were significantly enriched for four metabolic pathways (S2 Fig): “Fatty acid synthesis”, “Pyrimidine synthesis”, “Aminosugar metabolism” and “Purine synthesis”. These pathways are also retrieved in the studied DAR cluster (Fig 5). Similarly, DARs predicted for valproic acid were significantly enriched for 12 metabolic pathways (S2 Fig): “Pyrimidine synthesis”, “Aminosugar metabolism”, “Sphingolipid metabolism”, “NAD metabolism”, “Thiamine metabolism”, “Glycolysis/gluconeogenesis”, “Fructose and mannose metabolism”, “Lysine metabolism”, “Tryptophan metabolism”, “Urea cycle”, “Limonene and pinene degradation”, and “Hyaluronan metabolism” which is in line with results obtained from the graph-based analysis (Fig 4). Interestingly, pathway enrichment results are very different when artificial reactions (S6 Table) (*i.e.* sink, pool and extracellular exchange reactions) are kept in the list of DARs, suggesting that the list of DARs should be carefully processed before performing pathway enrichment (see S1 Appendix).

We observed that the fatty acid synthesis pathway was evidenced as significantly enriched from DARs for four molecules and compared how these four molecules were impacting this pathway (Fig 6) by visually comparing the metabolic footprint of each chemical on the fatty acid synthesis pathway. Amiodarone and allopurinol affected the same reactions (Fig 6 A and Fig 6 C), which suggested that they share similar mMoAs regarding fatty acid synthesis. On the other hand, rifampicin and sulindac seemed to impact another part of the fatty acid synthesis pathway (Fig 6 B and Fig 6 D). These observations showed that, although pathway enrichment can provide information on which general pathways are impacted by a chemical, it is not sufficient to evidence more precise modulated functions or to differentiate between potential different mechanisms of action. Even if these four compounds are identified as being able to induce liver steatosis, underlying MoAs might differ [51]. It is then interesting to see how our approach might allow in-depth understanding of MoAs.

**Fig 6.**
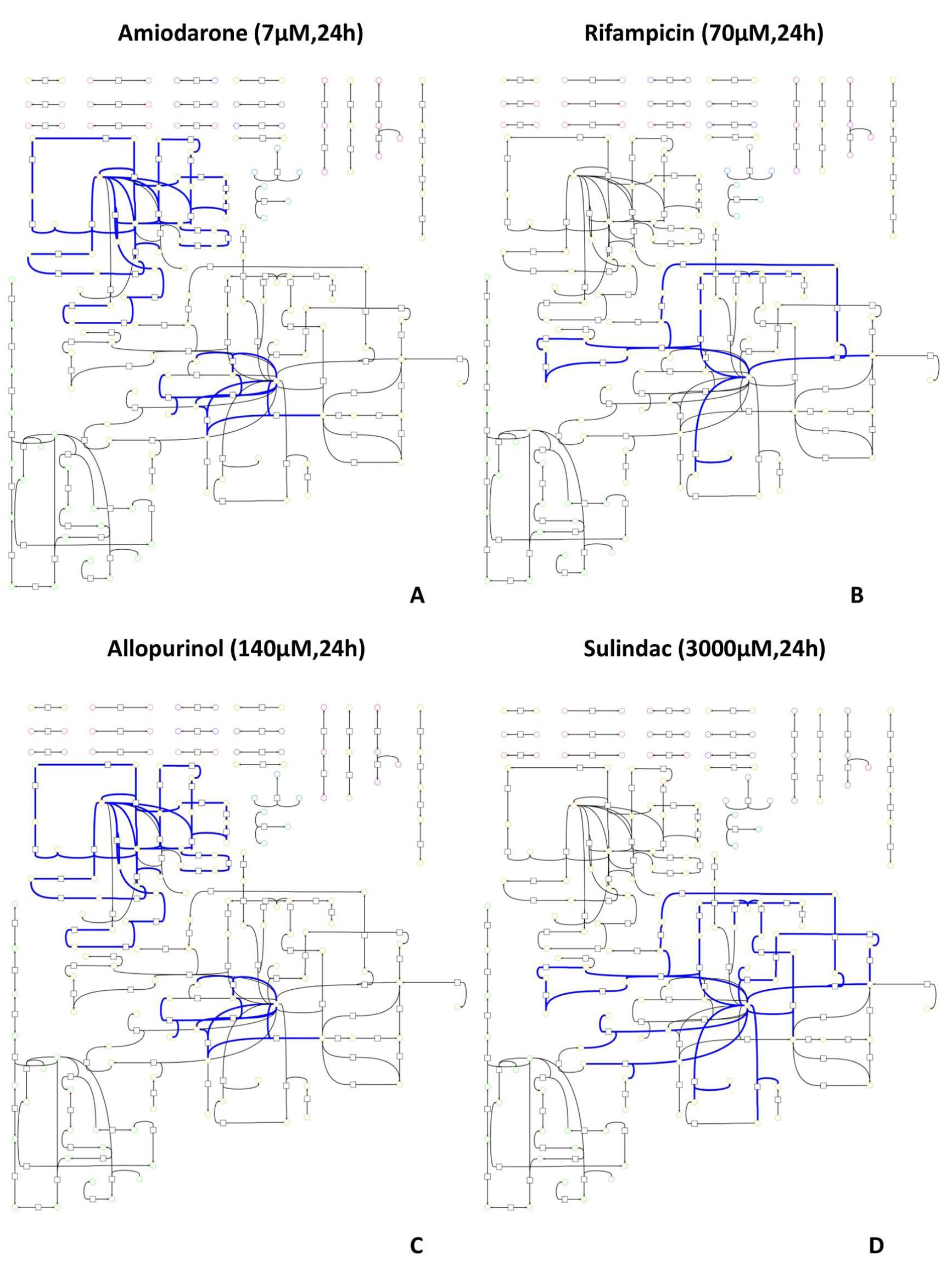
Comparison of the metabolic impact of four molecules with fatty-acid synthesis pathway significantly enriched. Each metabolic graph represents which fatty acid pathway reactions are differentially activated (in blue) after *in vitro* exposure to amiodarone (Fig 6 A), sulindac (Fig 6 B), allopurinol (Fig 6 C), and rifampicin (Fig 6 D). Nodes represented by a square are metabolic reactions and nodes represented by a circle are metabolites. DARs links have been highlighted in blue to identify which part of the pathway is perturbed by the molecule.

### Metabolic distance-based clustering and subnetwork extraction for mMoA interpretation

To go beyond pathway enrichment-based analysis, we propose a clustering approach based on the metabolic distance in the network, which ensures the identification of groups of functionally interdependent modulated reactions. These groups can involve reactions belonging to different pathways, therefore enabling the study of the mMoA as a continuous phenomenon instead of a binary (*i.e.*, enriched/not enriched pathway) phenomenon. However, hierarchical clustering suffers from some limitations. Indeed, hierarchical clustering is not designed to capture communities (*i.e.*, groups of densely interconnected elements in a graph) and tends to consider reactions implicated at the ends of linear cascade as distant, therefore not biochemically related, even if according to the network topology these reactions are biochemically related and highly dependent. Community detection approaches could be an answer to some of these limitations but their performance may be affected by the particular topological properties of large networks [52] such as GSMNs, therefore requiring a thorough evaluation to find the most effective method.

To add metabolic context and functional information to the lists of DARs and find connected subnetworks that include clustered DARs, we extracted minimal subnetworks with an algorithm based on the Steiner minimal tree problem. This algorithm is well adapted to our principal objective of visualizing human-readable subnetworks because it will find the smallest subnetwork connecting all the nodes of interest (*i.e.*, DARs) while adding as few nodes as possible for connectivity. However, the Steiner minimal tree problem is computationally hard (proven to be NP-Complete [53]). The metric closure graph [54] (or shortest distance graph) approximation was thus used. While employing an approximation is necessary, it introduces an arbitrary decision-making process within the algorithm, leading to the selection of a single alternative minimal Steiner tree among various possibilities. This choice, although necessary to extract a minimal subnetwork representative of the mMoA, could miss biologically interesting connections between DARs. The drawback of favoring the obtention of small graphs is that some interesting alternatives could be omitted. Another option could therefore be the k-shortest paths algorithm, which would make it possible to find these alternatives, at the expense of readability. Hence, regarding subnetwork extraction, method choice is guided by finding the right trade-off between readability and exhaustivity.

Because metabolic disruptions can impact a wide range of cellular functions spanning several metabolic pathways, the clustering and subnetwork extraction methods are key to connect and analyze disruptions. Interestingly, in the subnetwork presented in Fig 4, most of the reactions added by the subnetwork extraction algorithm are transport reactions, which highlight how the subnetwork extraction algorithm can help to better understand the mMoA of the molecule as a continuous phenomenon spanning several pathways and compartments rather than localized and disconnected perturbations. The graph-based analysis developed in this strategy could be used with other approaches able to identify perturbed reactions between two conditions, such as MOOMIN.

### mMoA analysis of PHH exposed to amiodarone or valproic acid

. The perturbations predicted by our workflow and visualized in Fig 4 and Fig 5 for Amiodarone and valproic acid are mainly associated with -oses and lipid metabolism, which is line with Amiodarone and valproic acid being known to induce hepatotoxicity via the occurrence of steatosis [41,42]. Valproic acid is also widely known for its impact on mitochondrial function [42] and is a histone de-acetylase inhibitor [55], a class of molecules known to impair many cell functions: the wide diversity of pathways subparts identified in the mMoA of valproic acid tend to support this mechanism (Fig 4B).

To study the metabolic impact of valproic acid on PHH after 24h exposure, we focused on visualizing cluster 2 (Fig 4), which is the cluster with the highest DARs coverage for this condition. We observed that DARs from this cluster were associated with 12 pathways. We were also able to identify another group of four up-activated reactions that is associated with lysine degradation in mitochondria, a phenomenon associated with mitochondrial homeostasis disruption in mice [56]. This phenomenon could also be due to a decrease in the L-carnitine pool associated with valproic acid exposure [57,58] that could trigger a compensating mechanism to restore L-carnitine which requires lysine as a substrate. Finally, a group of eight reactions predicted as up-activated is associated with the metabolism of tryptophan. Interestingly, an increase in the metabolism of tryptophan and kynurenine has already been reported as a potential effect of valproic acid in rats [59], and the valproic acid-induced increased conversion of tryptophan to nicotinamide has been observed in rats [60].

The impact of amiodarone on the fatty acid synthesis pathway predicted by our workflow (Fig 5) has already been observed in HepaRG cell cultures [51]. It is suggested that the increase in fatty acid synthesis could be due to the activation of SREBP1, which is a transcription factor mediating the expression of several important genes for *de novo* lipogenesis. One of these genes, FASN, encodes the fatty-acid synthase, which our workflow predicted as DAR in the treated state (Fig 5 A). Interestingly, SREBP1 and FASN were not identified as differentially expressed in amiodarone’s transcriptomic data, suggesting that our workflow is able to retrieve mechanistic information from lowly informative transcriptomic data. A similar increase of *de novo* lipogenesis on 3T3L1 adipocytes has also been reported [61] and is linked to an increase in palmitate production, which is in line with the predicted up-activation of the fatty-acyl-ACP hydrolase hydrolyzing palmitoyl-ACP to palmitate (Fig 5). In the subnetwork presented in Fig 5, only two reactions have been added by the subnetwork extraction algorithm, suggesting that DARs identified in this cluster are closely located in the metabolic network, therefore functionally interdependent. Moreover, the visualized subnetwork accounts for 74% of predicted DARs for amiodarone. Since the majority of DARs in the visualized subnetwork (Fig 5) are associated with fatty acid synthesis, predictions from our workflow suggest that amiodarone mMoA is mostly related to this pathway and quite localized in the metabolic network. This mechanistic observation would not have been possible with the initial transcriptomic data analysis that yielded only two DEGs for PHH exposed 24h at 7µM amiodarone, thus emphasizing the capacity of our workflow to extract mechanistic information from condition-specific metabolic networks constructed with lowly informative transcriptomic data.

## 4 Conclusion

In this study, we proposed a strategy dealing with the increased complexity of performing partial enumeration during condition-specific modeling by identifying DARs and visualizing the chemical’s mMoA with a graph-based approach. The proposed strategy starts by integrating metabolic gene expression data to a GSMN to construct sets of condition-specific metabolic networks. These sets of conditions-specific metabolic networks are then analyzed through an original statistical approach to identify reactions that are likely to be differentially activated between the two compared conditions. To go further on the mechanistic understanding, we implemented a graph-based network analysis step to extract and visualize subnetworks corresponding to metabolic functional and connected groups of DARs. Finally, we showed that the strategy that we developed succeeded in retrieving parts of the mMoA described in the literature for two well-known hepatotoxic molecules (amiodarone and valproic acid), even when only few genes were significantly disrupted.

The presented strategy provides a way to take advantage of the increased robustness obtained with condition-specific enumerated networks while keeping the analysis doable and human readable, allowing a better and deeper understanding of chemicals’ mMoA. This strategy can be divided in several independent parts: data integration, condition-specific modelling and graph-based analysis, so that it can be combined with other condition-specific modelling approaches. This comprehensive metabolic network exploration and visualization comes at the expense of the computation time, which could make it challenging to apply this strategy in large-scale safety assessment studies and would require development in parallel of complementary methods providing a more global assessment of the metabolic impact of a compound. These complementary developments could be plugged into the current workflow quite easily thanks to its flexibility. Although we focused on studying the metabolic impact of chemicals on PHH, our workflow could also be used to model the metabolic impact of chemicals on different cellular types from different tissues, therefore paving the way for a more precise understanding of how a chemical impacts the metabolism of many organs.

## 5 Methods

### Transcriptomic data processing

Transcriptomic data used in this study were obtained from the Open TG-GATEs [21] database. For this study, we selected *in vitro* data generated on PHH. Raw transcriptomic data were downloaded as CEL files from https://dbarchive.biosciencedbc.jp/en/open-tggates/download.html. CEL files were read with the affy [62] package and the resulting dataset was normalized with the robust multiarray average method [63] from the limma R package. Information about the different PHH cell batches used in the different experiments was obtained from the authors (S5 Table) and was used to correct the batch effect with Combat [64], available in the SVA package. We annotated the probes with the annotation database corresponding to the Affymetrix HG133U Plus 2 chips available in the hgu133plus2.db package [65] and the AnnotationDbi package [66]. Where several different probes mapped to the same gene, we selected the probe with the highest standard deviation of expression values in the dataset. Finally, to match DEXOM [34] requirements, we categorized transcriptomic data with Barcode [67–69]. We used the z-scores output available from the barcode package and applied a 25/75 percentiles cutoff to obtain a list of highly and lowly expressed genes. Genes below the 25^th^ percentile were considered as lowly expressed genes and assigned a -1 value, while genes above the 75^th^ percentile were considered as highly expressed genes and assigned a +1 value. Genes between the 25^th^ and 75^th^ percentiles were not considered (*i.e.*, 0 value) and therefore did not have any impact on the modeling process.

### DEG identification

Lists of DEGs were obtained from the ToxicoDB [20] database, which provides pre-processed differential gene expression data from several databases, including Open TG-GATEs. A jupyter notebook querying ToxicoDB API was developed to automate the process. First, it downloads the ToxicoDB compounds json file and iterates over the list of compounds obtained from this file to query the ToxicoDB API to get analyzed data associated with each compound. The retrieved data were then filtered by selecting genes from the Open TG-GATEs human study with an absolute value of log2(FC) larger than 0.26 and a q-value less than 0.05, from samples subjected to a “high” dose (three doses were investigated and available for this dataset: low, middle, and high) over 24 hours. The filtered DEGs list for each compound was then stored in a .tsv file.

### Metabolic model preparation

We used the Recon2.2 [26] (downloaded from https://www.ebi.ac.uk/biomodels/MODEL1603150001) GSMN as the initial framework for the modeling and network analysis steps. This GSMN is composed of 5324 metabolites, 7785 reactions, and 1675 genes. We modified the model biomass reaction, to account only for the precursors required for cell maintenance but not replication, in order to better represent the fact that PHH are differentiated cells with a short lifespan and do not proliferate under classical culture conditions [70]. To do so, we set the stoichiometric coefficient of the DNA precursor in the biomass reaction to zero and applied a correction coefficient (S1 Appendix) to other metabolites to keep the reaction balanced. Next, we forced the lower bound of the modified biomass reaction, which is now representing the cost of maintaining the cell viability, to a value of 1 instead of 0 to ensure that the generated models will be able to maintain cell viability (*i.e.*, have a non-zero flux through the maintenance reaction). Exchange reactions were left unconstrained as we did not have information on cell medium composition.

### Condition-specific modeling with partial enumeration

We integrated the categorized transcriptomic data into the GSMN through GPRs to define an *a priori* set of active and inactive reactions. A highly expressed gene will be given the value of 1 whereas a lowly expressed gene will be given the value of -1. Genes are associated with reactions by the GPRs. GPRs are Boolean rules that indicate which genes are required to produce a specific enzyme that catalyzes one or more reactions. When a reaction has an AND GPR, the algorithm will annotate the reaction as active if the minimal value of the categorized GPR’s gene expression values equal 1 (*i.e.*, all genes of the GPR are highly expressed) and inactive otherwise, such as:

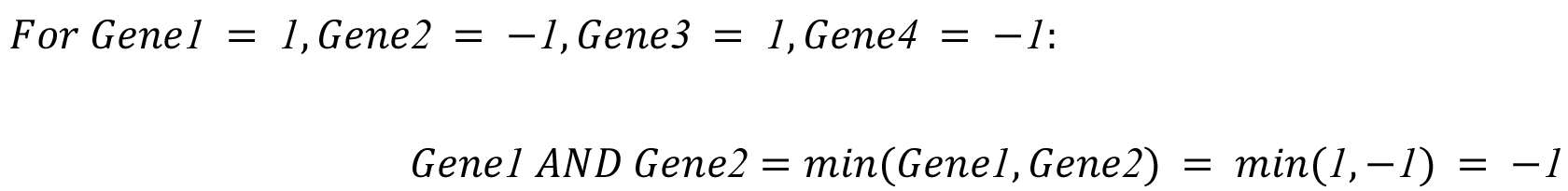

In this case, if *Gene1* is highly expressed *AND Gene2* is lowly expressed, the reaction is considered inactive.

When a reaction has an OR GPR, the algorithm will annotate the reaction as active if the maximal value of the categorized GPR’s gene expression values equal one (*i.e.*, at least one gene of the GPR is highly expressed) and inactive otherwise, such as:

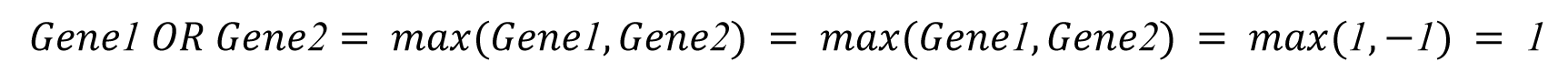

In this case, if *Gene1* or *Gene2* is highly expressed, the reaction is considered active. These rules can be applied to more complex GPRs such as:

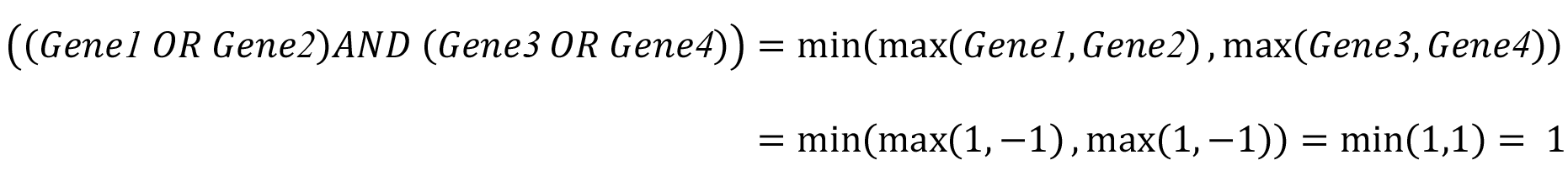

In this case, if *Gene1* or *Gene2* is highly expressed and *Gene3* or *Gene4* is highly expressed, the reaction is considered active.

A reaction will be identified as *a priori* active if the resulting value equals 1 and as *a priori* inactive if the resulting value equals -1. At the end of this process, we identified a list of *a priori* active/inactive reactions for each sample that was then provided to DEXOM. DEXOM is a constraint-based modeling algorithm based on iMAT [29,30] whose objective is to find a steady state distribution of flux that maximizes the number of reactions whose flux is consistent with transcriptomic data levels [30].

To extract a representative set of solutions from the whole solution space, we adapted the partial enumeration from DEXOM and used the python implementation available in dexom-python (https://github.com/MetExplore/dexom- <u>python</u>). First, we applied a full Reaction-enum strategy (see [34] for the complete description). This method iterates over all the reactions of the network, blocking each of them successively and solving the resulting mixed integer linear problem. Next, we applied the DEXOM Diversity-enum strategy, starting with a random solution picked per range of Reaction-enum solutions as a starting point. Diversity-enum aims to find new solutions that are gradually more different from the starting solution, allowing one to explore the limits of the solutions’ space. To reduce the computational costs of the original DEXOM partial enumeration approach, we reduced the number of starting solutions to a set of 1% of the solutions obtained with the Reaction-enum approach. We used an adapted version of systematic sampling (*i.e.*, one random solution picked per batch of solutions, S1 appendix for details) to generate a starting set representative of the complete set of Reaction-enum solutions. Finally, all the optimal solutions for a sample were grouped in a single tabulation-separated file. The list of parameters used for running the DEXOM algorithm can be found in S1 Appendix.

### Identification of differentially activated reactions

For each reaction, the predicted activation status across all optimal condition-specific metabolic networks enumerated by the DEXOM strategy (Table 1) is stored in a numeric vector. Therefore, for each condition, the output of the DEXOM enumeration is a matrix of binary vectors where columns correspond to reactions and rows correspond to enumerated optimal solutions (Table 3).

**Table 3.** DEXOM output example for one condition. Alternative solutions obtained with partial enumeration are stored as binary vectors.

| *DEXOM solution* | **Reaction 1** | **Reaction 2** | **Reaction 3** | ... | **Reaction N-1** | **Reaction N** |
| --- | --- | --- | --- | --- | --- | --- |
| *Solution 1* | 0 | 0 | 0 | ... | 1 | 1 |
| .... | ... | ... | ... | ... | ... | ... |
| *Solution N* | 1 | 1 | 0 | ... | 0 | 1 |

For each reaction and each condition, we can compute an activation frequency value *f*, as the number of solutions in which the reaction is predicted to be active, over the total number of solutions. To compare the activation frequency values of two conditions, we introduce a metric called R2.

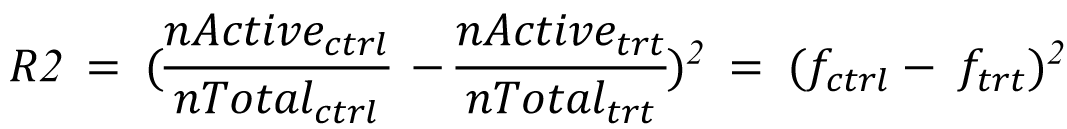

Where *nActive* is the number of times the reaction is predicted active and *nTotal* is the total number of solutions in the condition (*i.e.*, *nActive*+*nInactive*). *f*_ctrl_ is the activation frequency of a reaction under the control condition and *f*_trt_ is the activation frequency of the same reaction under the treated condition.

We considered reactions as differentially activated if the R2 value was higher than 0.2 when comparing treated *vs* non-treated conditions. This is a rather conservative threshold since it means that to be considered as differentially activated, a reaction needs to be at least almost twice (80%) more/less active in the treated condition compared to the non-treated condition.

### Baseline noise calculation

The absence of gene expression information for some reactions, and therefore the absence of constraints on those reactions when using the DEXOM algorithm, leads to uncertainty (or “noise”) in the reaction activation frequency prediction. Indeed, in the absence of transcriptomic data on reactions, DEXOM can predict these reactions as active or inactive without any impact on the optimality score. This uncertainty could lead to biased conclusions when comparing two conditions. To avoid considering reactions that are loosely constrained as DAR, we implemented a method that aims at estimating the baseline noise associated with each reaction in the network. The noise refers to the variation of the activation frequency of each reaction between pairs of control conditions, which is due to the fact that the predicted activity of the reaction does not impact the consistency with the data. Because several control conditions and chemical exposure times have been used in this dataset, we computed the baseline noise for each reaction between control conditions at a specified time and a specified vehicle. Replicates were pooled for each molecule. Practically, for each reaction, we computed the R2 between all pairs of control conditions for a given vehicle at a given exposure time and calculated the median of all R2s for each reaction. Therefore, we obtained for each reaction a baseline noise estimation that is the median of all pairwise comparisons between selected control conditions.

Then we filtered out DARs with an R2 between two conditions of interest (*e.g.*, control *vs* treatment) that was less than two times the noise estimate for this reaction.

### Metabolic reaction graph construction

GSMNs, which are large and complex metabolic networks, contain hub nodes (*i.e.*, nodes connected to many other nodes). These hub nodes correspond mainly to cofactor compounds, *i.e.,* metabolites required in many biochemical reactions (ATP, ADP, NADPH, etc.) therefore creating connections between many reactions, although these connections are often not biologically meaningful. This property of GSMNs is a challenge [71] for network analysis approaches based on graph properties and topology. To solve this issue, we identified a list of “side compounds” corresponding to the main hub nodes of the network (S3 Table) that will be removed during the metabolic graph construction. To improve the performance of the graph theory methods, we also identified a list of reactions (S6 Table) to be excluded during the reaction metabolic graph construction. These reactions are blocked reactions (*i.e.*, reactions that cannot carry a flux in the GSMN), reactions always inactive in our cellular model, pool, sink, and exchange reactions. We used the metabolic reaction graph representation, where nodes are reactions and edges connect two reactions if the product of one is the substrate of the other. Metabolic reaction graphs were constructed according to the parameters defined previously before applying any graph-based methods such as the distance matrix calculation and subnetwork extractions.

### Reaction distance matrix calculation

We computed the pairwise distance between reactions of interest by computing all the shortest paths between the nodes of the network corresponding to these reactions, resulting in a reaction pairwise distance matrix. To that purpose, we implemented a java app using the Met4j metabolic network toolbox (https://forgemia.inra.fr/metexplore/met4j). This app takes as input the GSMN (*i.e.*, Recon2.2) SBML file, a list of side compounds, a list of reactions to exclude, and a list of reactions between which the distances will be computed (*e.g.*, DARs in our case).

After loading the SBML file, the GSMN is pruned by removing side compounds and reactions to exclude. Then the JGraphT [72] ManytoMany shortest path implementation [73] is used. This implementation has been preferred over more common implementations such as Dijkstra [74] or Floyd–Warshall [75] due to its performance and its ability to take a specific list of reactions as input. The distance matrix is then saved in a comma separated file.

### DARs clustering

For each DARs distance matrix computed with Met4J, we performed a hierarchical clustering with the Ward algorithm implemented in the SciPy python module. To control the size of clusters, we visualized each clustering result with a dendrogram representation and manually determined the number of clusters. Clusters were then obtained with the cutree function.

### Subnetwork extraction

For each cluster of DARs, we extracted a minimal subnetwork with the minimal Steiner tree algorithm implemented in Met4J. This algorithm searches for a minimum spanning tree that contains the set of DARs and a minimum set of nodes from the GSMN to connect them according to the initial network topology.

### Subnetwork visualization

We visualized the extracted subnetworks with MetExploreViz [40]. MetExploreViz is a JavaScript library dedicated to visualization for GSMNs that is integrated within the MetExplore [76] webserver. The list of metabolic reactions contained in the subnetwork is first mapped on the selected GSMN with MetExplore (https://metexplore.toulouse.inrae.fr/index.html/). From this mapping, we can then visualize and prune (*i.e.*, remove side compounds) the bipartite metabolic graph interactively with MetExploreViz. Two visualizations were done for each of the subnetworks. One with mapped up-activated DARs colored in red and down-activated DARs colored in green (Fig 4 A and 4 B), and a second one with reaction links colored according to the pathway associated with the reaction (Fig 4 B and 5 B). Interactive visualizations were saved online and are accessible via links provided in supplementary materials.

### Workflow implementation

We partitioned the workflow into three jupyter notebooks with a common properties file. The first notebook, named “partial_enumeration.ipynb,” takes as input barcode z-scores and generates batch files to execute the partial enumeration protocol on a SLURM computing cluster. Of note, these batch files can also be executed directly on a standard Linux (*i.e.*, Ubuntu, Debian, etc.) operating system. The second notebook, named “dars_calculation.ipynb,” takes as input the partial-enumeration results from the previous notebook and computes baseline noise and DARs. Finally, the third notebook, named “analysis.ipynb,” handles the network analysis step of our workflow, which includes computing the distance matrix for each list of DARs, hierarchical clustering, and extracting subnetworks with Met4J.

Python library code and jupyter notebooks are available on GitLab: https://forgemia.inra.fr/metexplore/MANA

### Computing environment

Condition-specific modeling and partial enumeration requires CPLEX v20.10 to be installed on your system. Batch files generated by the “partial_enumeration.ipynb” notebook are designed to be launched on a SLURM computing grid but can also be launched on a standard Linux operating system. A local version of dexom-python is also required and can be cloned from the dexom-python repository (https://github.com/MetExplore/dexom-python). Finally, the network analysis workflow calling Met4J apps requires at least Java 11 to be installed on your machine.

## Declarations

### Ethics approval and consent to participate

Not applicable

## Consent for publication

Not applicable

## Availability of data and materials

All data used in this study were obtained from the Open TG-GATEs [21] database. Raw transcriptomic data were downloaded as CEL files from https://dbarchive.biosciencedbc.jp/en/open-tggates/download.html.

## Competing interests

## Authors’s contributions

LF, OP, AR, RG, FJ and NP designed the research; LF analysed the data; LF, AO, JCG, MS, CF FJ and NP contributed to the development of the methodology (including development and implementation of statistical, modelling and visualisation methods). AR, OP, BF, FJ and NP supervised the research; all authors read and approved the final manuscript.

## Acknowledgements

We thank Dr. Hiroshi Yamada for providing supplementary data regarding the cell batches for the assays from the Open TG-GATEs database. We also thank Dr. Pablo Rodriguez-Mier for his expertise on how to tailor DEXOM to our needs and the discussions regarding condition-specific modeling and partial enumeration. Finally, we thank Dr. Gladys Ouedraogo for proofreading the manuscript and discussions throughout the project, and Dr. Bathilde Ambroise for the very interesting discussions we had during the very first phase of this work.

## Supplementary materials table of contents

**S1 Appendix Supplementary information.** Pdf including: Correction coefficient formula, DAR specificity ratio formula, DEXOM parameters, Systematic sampling.

**S1 Table DAR filtered out by the baseline noise filter.** This table lists DAR having a R2 that is less than two times the noise estimated for this reaction, therefore filtered out by the baseline noise filter.

**S2 Table DAR metrics.** This table contains several metrics which aim at understanding how perturbed reactions are commonly perturbed between the eight studied chemicals. This table lists size, specificity ratio and average percentage of DARs intersection size.

**S3 Table Side compounds identifiers.** This table lists the sides compounds used in the network analysis stages.

**S4 Table DAR subnetwork coverage.** This table lists the DAR subnetwork coverage for each identified subnetwork for amiodarone and valproic acid.

**S5 Table DEG lists size obtained from the ToxicoDB database.** This table summarize key metadata for each compound such as the dose, exposure time, number of replicates, cell type, cell batch, control vehicle, DEG signature size and metabolic DEG signature size. Were considered differentially expressed, genes having a log2(abs(FC)) above 0.26 and a FDR corrected p-value below 0.05.

**S6 Table Excluded reactions identifiers.** This table lists the reactions that have been excluded in the network analysis stages.

**S1 Fig Pathway enrichment performed on valproic acid DEGs with ReactomePA.** Pathway over-representation analysis performed with a Fisher Exact test, p-values corrected with the Benjamini-Hochberg method on Reactome 2022 pathways (genes with log2(absFC)) > 0.26 and FDR-corrected p-values < 0.05 were considered as DEG)

**S2 Fig Pathway enrichment performed on DARs, after removing blocked and artificial reactions.** Pathway over-representation analysis was performed on Recon2.2 metabolic pathways for DARs predicted for the eight selected hepatotoxic molecules? DARs were filtered to remove blocked and artificial reactions. P-values were computed using a Fisher Exact test, with a Benjamini-Hochberg correction.

**S3 Fig Pathway enrichment performed on DARs.** Pathway over-representation analysis was performed on Recon2.2 metabolic pathways for DARs predicted for the eight selected hepatotoxic molecules? DARs were filtered to remove blocked and artificial reactions. P-values were computed using a Fisher Exact test, with a Benjamini-Hochberg correction. S4 Fig Biclustered heatmap of the pairwise reaction distance matrix of DARs predicted for amiodarone (A) and valproic acid (B). Hierarchical biclustering was performed using the Ward algorithm on the pairwise reaction distance matrix for DARs predicted for amiodarone and valproic acid. The resulting biclustering was visualized as a heatmap computed with the Pheatmap R package. The color scale depicts the distance between two reactions. The distance ranges between zero (cells colored in blue) to eight for amiodarone (A) and 14 for valproic acid (B) (cells colored in red). Two main clusters (C1 and C2) were identified for amiodarone and 4 clusters (C1, C2, C3, and C4) were identified for valproic acid.

