## Supplementary material for "A strategy to detect metabolic changes induced by exposure to chemicals from large sets of condition-specific metabolic models computed with enumeration techniques": S1 Appendix

### Interactive visualizations with MetExploreViz:

Fig 4A: https://metexplore.toulouse.inrae.fr/userFiles/metExploreViz/index.html?dir=/72ff7fdc7031b880ef4f3532134aa326/networkSaved_373423088

Fig 4B: https://metexplore.toulouse.inrae.fr/userFiles/metExploreViz/index.html?dir=/72ff7fdc7031b880ef4f3532134aa326/networkSaved_725935955

Fig 5A: https://metexplore.toulouse.inrae.fr/userFiles/metExploreViz/index.html?dir=/72ff7fdc7031b880ef4f3532134aa326/networkSaved_292937465

Fig 5B: https://metexplore.toulouse.inrae.fr/userFiles/metExploreViz/index.html?dir=/72ff7fdc7031b880ef4f3532134aa326/networkSaved_1994092833

### Correction coefficient:

### To maintain the biomass reaction balanced after setting the stoichiometry coefficient for the biomass_DNA reactant to zero, we applied the following correction coefficient to the stochiometric coefficient for all other reactants of the biomass reaction:

##

$$\boldsymbol{corCoef =}\frac{\boldsymbol{1}}{\boldsymbol{1+}\boldsymbol{S}_{\boldsymbol{DNA}}}$$

Where $S_{DNA}$ refers to the stoichiometric coefficient for the biomass_DNA reactant in the biomass reaction initially defined in Recon2.2 [1]. We applied this correction coefficient to all other metabolites (i.e biomass_lipid, biomass_carbohydrate, biomass_other, biomass_RNA and biomass_protein) of the Recon2.2 biomass reaction

### DAR specificity ratio:

### To evaluate the degree of specificity of a Differentially Activated Reaction (DAR) signature, we compute the DAR specificity ratio. A DAR specificity ratio of 100% means that all the predicted DARs are retrieved only for the studied molecule, meaning that the metabolic Mechanism of Action (mMoA) of this molecule is unique compared to mMoA of other studied molecules.

##

$$\boldsymbol{DARspec}_{\boldsymbol{x}}\boldsymbol{=}\frac{\boldsymbol{nSpecificDARs}_{\boldsymbol{x}}}{\boldsymbol{nTotalDARs}_{\boldsymbol{x}}}$$

### With $\boldsymbol{nSpecificDARs}_{\boldsymbol{x}}$, the number of DARs retrieved only in the list of DARs predicted for the molecule X and $\boldsymbol{nTotalDARs}_{\boldsymbol{x}}$, the total number of DARs predicted for the molecule X.

### DEXOM parameters:

The choice of parameters values is based on a compromise between the quality of returned solutions and the solving time.

Indeed, we can compare the optimality score achieved by the algorithm to the theoretical optimality score of the constructed mixed integer linear problem which is the maximum adequacy score between the network topology of the GSMN and the list of active/inactive reactions obtained from the binarization of transcriptomic data.

We selected the following parameters:

• eps = 1e^-2^

• thr = 1e^-5^

• tlim = 600s

• mipgaptol = 1e^-3^

eps and thr are the thresholds above which a reaction is considered active. eps represent, the activation threshold of activated reactions (according to transcriptomic data) and thr represents, the activation threshold for unweighted reactions (according to transcriptomic data). tlim is the parameter defining the maximum time allowed to solve a mixed integer linear problem. The mipgaptol parameter defines the maximal allowed gap between the theoretical maximal adequacy score and the adequacy score of the optimized solution allowing us to consider the solution as optimal. The trade-off between quality and running time described above is mainly affected by tlim and mipgaptol. Therefore, these two parameters might need to be carefully adjusted according to the complexity of the problem and the available computational resources.

### Adapted version of Systematic sampling:

Systematic sampling aims at picking random samples in a uniform fashion over all the dataset [2]. Here, the dataset consists of solutions enumerated with the Reaction-Enum partial enumeration method. We implemented this systematic sampling method because in Recon2.2, reactions are sorted in alphabetical order by default and some reactions names are informative of their function (*e.g.* EX_R… refers to Exchange reactions). Since the aim of Reaction-Enum is to iteratively block each reaction and if possible, find an optimal solution with this reaction blocked, the order of solutions returned by Reaction-Enum is impacted by this reaction naming bias.

Therefore, to avoid picking an imbalanced set of starting solutions (*i.e.* a set of solutions not representative of the diversity of possible solutions computed by Reaction-Enum), we implemented a systematic sampling approach.

First, we compute the number of batches according to the size of the range defined at the beginning of the batch computation of the Reaction-Enum step. Then for each batch we pick a random solution in the corresponding range of Reaction-Enum solutions. This solution will then be used as starting point of a Diversity-Enum batch. As an example, in the case where we have 4500 solutions obtained by Reaction-Enum and a range of 100 solutions per batch, the starting solution for the first Diversity-Enum batch will be picked between the first and 99^th^ Reaction-Enum solutions, the starting solution for the second Diversity-Enum batch will be picked between the 100^th^ and the 199^th^ Reaction-Enum solutions and so on until we picked a starting solution for each batch, the last one being between the 4400^th^ and the 4500^th^ Reaction-Enum solutions

### Impact of DARs filtration on pathway enrichment results:

### Pathway over-representation analysis (ORA) is size sensitive [3,4]. Therefore, keeping reactions that are not functionally informative (*i.e.* sink, pool and exchange reactions) can lead to important differences in pathway ORA. For instance, when keeping such reactions, only one significantly enriched pathway (Transport, extracellular) was evidenced for valproic acid (S3 Fig), whereas 12 were identified when removing these reactions. These results suggest that DARs lists should be carefully processed in order to remove such reactions before performing an over-representation analysis. Interestingly, the graph-based analysis developed in this study is less impacted by keeping or removing these reactions since they are usually clustered together.
