## Supplementary figures and images for "A strategy to detect metabolic changes induced by exposure to chemicals from large sets of condition-specific metabolic models computed with enumeration techniques"

### S1 Fig

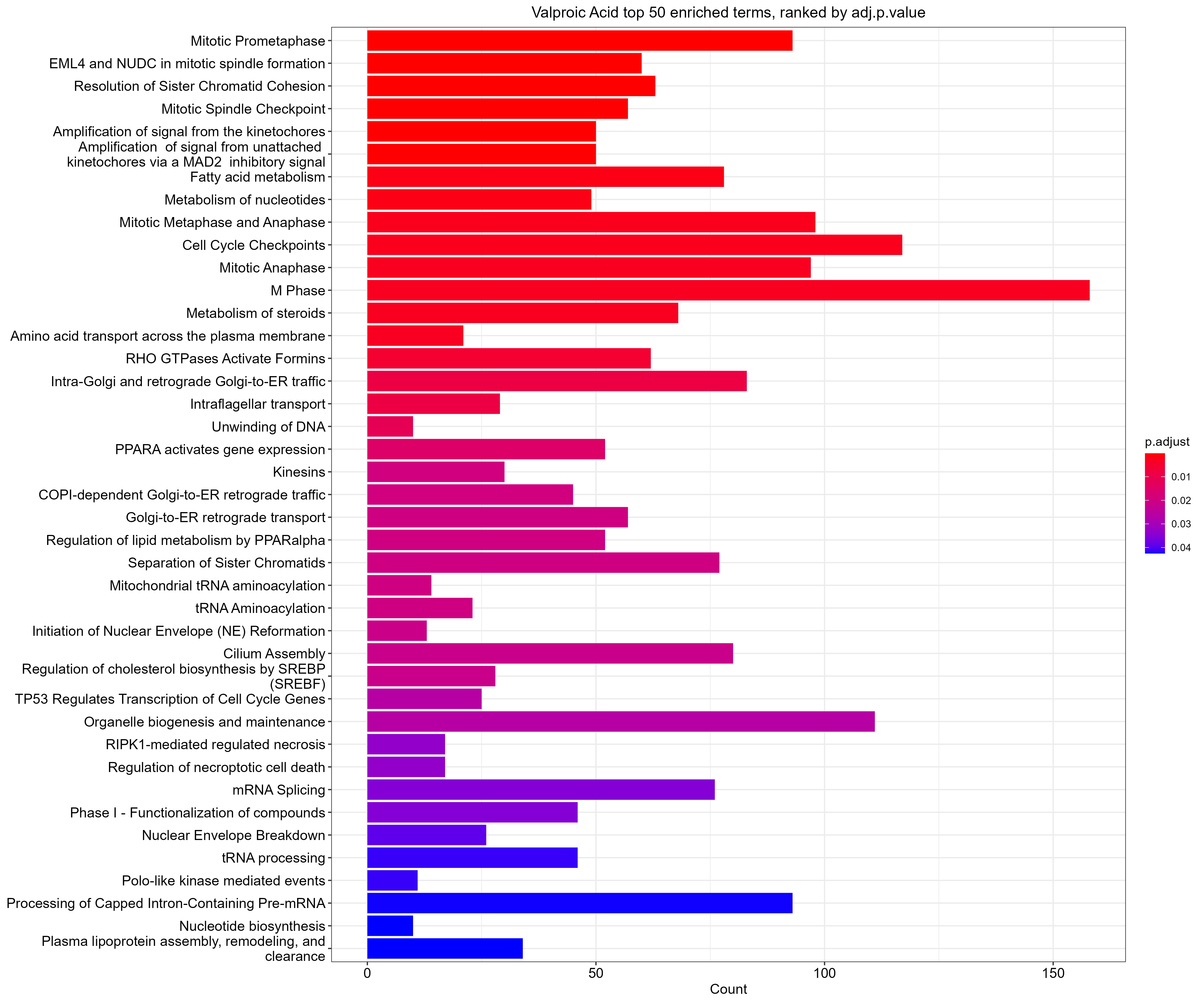

### S2 Fig

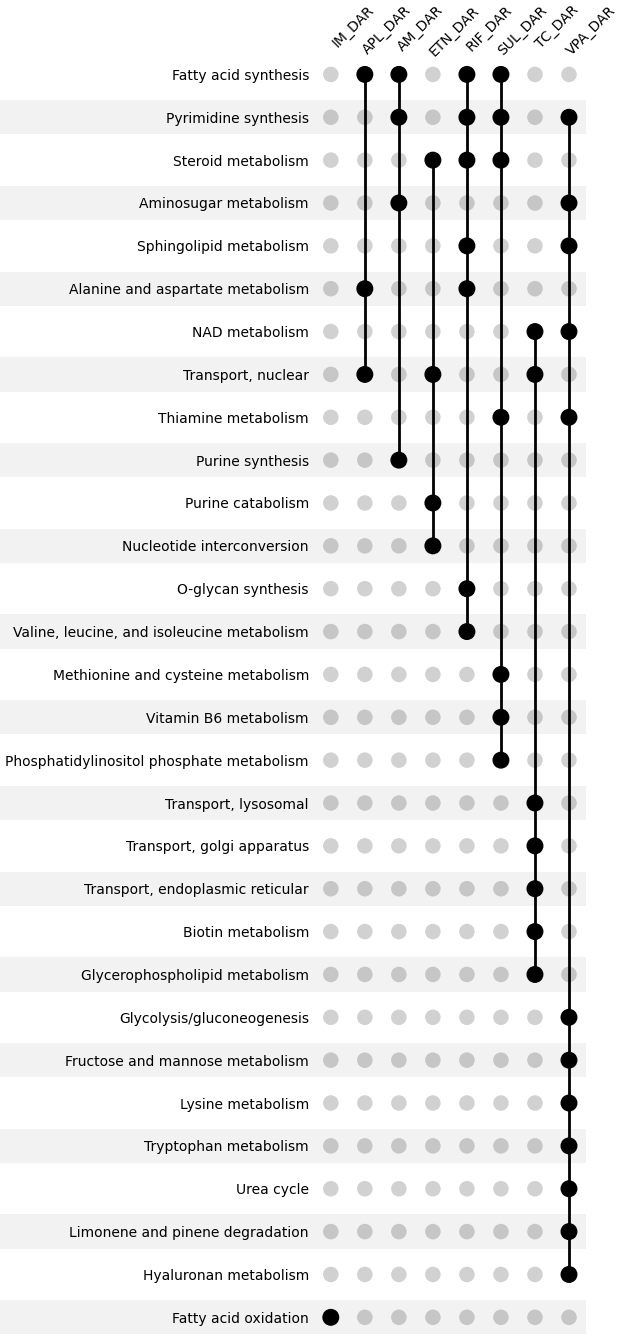

### S3 Fig

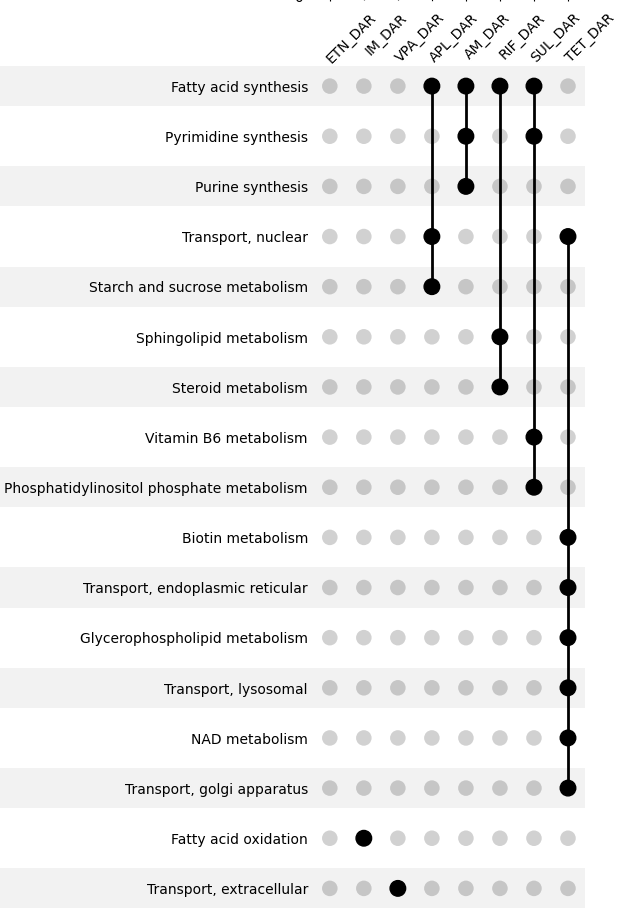

### S4 Fig

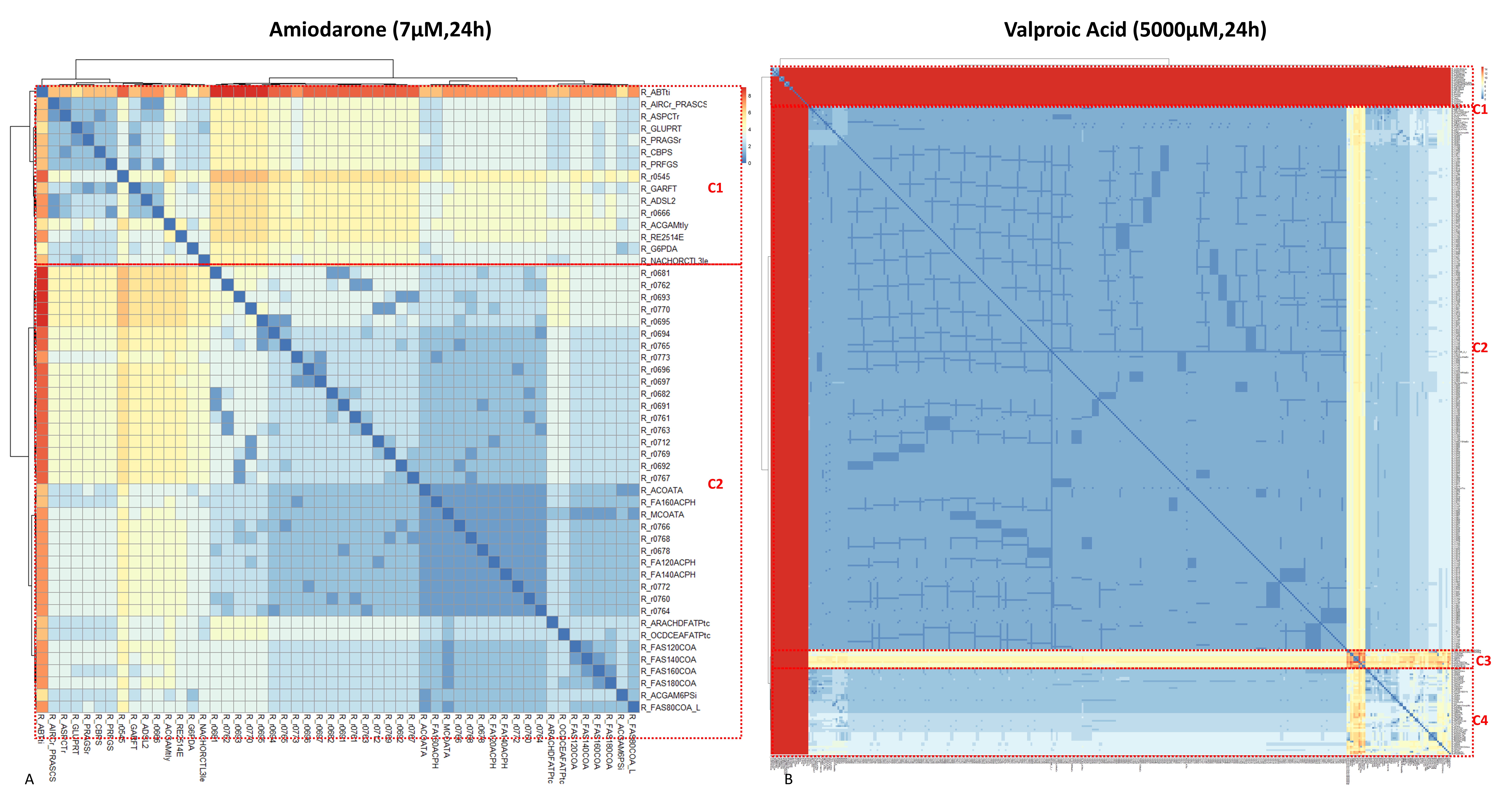
